# Fibroblast-specific protein-protein interactions for myocardial fibrosis from MetaCore network

**DOI:** 10.1101/2024.09.04.611186

**Authors:** Klaus M. Frahm, Ekaterina Kotelnikova, Oksana Kunduzova, Dima L. Shepelyansky

## Abstract

Myocardial fibrosis is a major pathologic disorder associated with a multitude of cardiovascular diseases (CVD). The pathogenesis is complex and encompasses multiple molecular pathways. Integration of fibrosis-associated genes into the global MetaCore network of protein-protein interactions (PPI) offers opportunities to identify PPI with functional and therapeutic significance. Here, we report the generation of a fibrosis-focused PPI network and identification of fibroblast-specific PPI driving reparative and reactive myocardial fibrosis. In TGFb-mediated fibroblast activation, PPI hubs predict new regulatory mechanisms for fibrosis-associated genes. We introduce an efficient Erdös barrage approach to suppress activation of a number of fibrosis-associated nodes in order to reverse fibrotic cascades. Our results suggest that PPI prediction model can offer network insights into fibrosis mechanisms and can complement future experimental efforts to counteract cardiac fibrosis.

## 1. Introduction

Myocardial fibrosis is a major pathologic disorder associated with a multitude of cardiovascular diseases (CVD) [1]. In the heart, tissue fibrotic remodeling is characterized by abnormal fibroblast activation and excessive extracellular matrix (ECM) protein accumulation [2]. Although a number of factors have been implicated in orchestrating the fibrotic response, tissue fibrosis is dominated by a central mediator: transforming growth factor-*β* (TGF-*β*) [3]. Sustained TGF-*β* production leads to a continuous cycle of growth factor signaling and deregulated matrix turnover [4]. However, despite intensive research, the biomolecules that orchestrate fibrosis are still poorly understood and as a result, effective strategies for limiting fibrosis are lacking [2,4]. Considering the complex heterogeneity of fibrosis, research strategy on a system-level understanding of the disease using mathematical modeling approaches is a driving force to dissect the complex processes involved in fibrotic disorders. Recently, we have reproduced the classic hallmarks of aberrant cardiac fibroblast activation leading to fibrosis and provided a powerful toolbox for characterization of cardiac fibroblast activation [5]. Although the pathogenesis of fibrotic remodeling has not been well identified, accumulated evidence suggests that multiple genes/proteins and their interactions play important roles in disease scenario [6].

Fibrotic remodeling is a complex process caused by genetic abnormalities that alter protein-protein interactions (PPI) [5,7]. In the heart, PPI interfaces represent a highly promising, although challenging, class of potential targets for therapeutic options. In fibrosis, PPI form signaling nodes and hubs that transmit pathophysiological cues along molecular networks to achieve an integrated biological output, thereby promoting fibrogenesis and fibrosis progression. Thus, pathway perturbation, through disruption of PPI critical for fibrosis, offers a novel and effective strategy for curtailing the transmission of pro-fibrotic signals. Deciphering of fibrosis-specific PPI would uncover new mechanisms of fibrotic signaling for therapeutic interrogation. Thus a mathematical analysis of the global PPI MetaCore network [8] can provide useful deep insights in the understanding of fibrosis progression processes.

In this work we use the PPI MetaCore network [8] to perform a mathematical modeling of fibrosis progression. The results reported in [5] determined the protein pro-fibrotic responses as a feedback on TGF protein stimulation. Thus these results [5] support the known fact that the TGF protein plays an important role in fibrosis tissue [3] and establish proteins with most positive and most negative response in cardiac fibroblasts.

A variety of biological applications of the MetaCore network are described in [9,10]. The general statistical properties of the MetaCore network are presented in [11]. The applications of Google matrix algorithms to the fibrosis PPI based on the MetaCore network are described in [12] using the fibrosis responses obtained in [5]. We note that the Google matrix algorithms [13–15] find a variety of useful applications in modern complex networks [16] including World Wide Web, Wikipedia, world trade etc. In our opinion an important advantage of the MetaCore network is that it presents a global network structure of PPI with about 40000 proteins and important molecules. This allows to perform a mathematical modeling of fibrosis progression over the whole network. Thus in this work we present such a modeling which assumes that at an initial stage there is a certain number of proteins which activate a fibrosis progression of other proteins via their network links to other proteins. These initial proteins are always marked as red network nodes and assumed to be always red permanently generating fibrosis progression (thus being always red or with fixed Ising spin up with *σ*_*i*_ = 1). Among the other proteins, we assume that there is an initial group of proteins which inhibit fibrosis progression being of fixed blue color (thus being always blue or with Ising spin down with *σ*_*i*_ = −1). The remaining other proteins (or network nodes) are assumed to be in neutral state of white color or spin being zero (*σ*_*i*_ = 0). The fibrosis progression is modeled by the asynchronous Monte Carlo process when at each step a spin of a given node (which is not fixed) is determined as the total spin sign of other nodes linked to it with certain weights of Markov chain transitions. In a certain sense this rule corresponds to a situation when a person takes an average opinion of his network friends linked to him. After many of such Monte Carlo steps the system converges to a steady-state when all network spins obtain a fixed polarization being up (red or fibrosis activated) or down (blue or healthy protected) and some of the nodes may remain at zero neutral spin state (white) being practically disconnected from the red or blue network nodes.

We note that the above Monte Carlo steps describe a process of opinion formation and the global vote of nodes (society members) between red and blue options. For complex social networks [16] there are numerous studies of opinion formation, reviewed in [17]. Various interesting features of voter models and opinion formation on networks had been obtained and described in [17,19–24]. Recently it was argued that the opinion formation on the world trade network can be linked with country preference to trade in one or another currency (e.g. US dollar or hypothetical BRICS currency) [25]. The important new element appeared in [25] is that opinion of certain network nodes (countries) is considered to be fixed since it is assumed that they prefer to trade always with fixed currency of USD or BRICS. A further development of this approach with nodes of fixed opinion is done in [26] for Wikipedia networks when initially there are two groups of nodes of fixed red and blue opinions but all other nodes have neutral state (white opinion or spin zero) and their fixed opinion emerges in the result of asynchronous Monte Carlo process. This process is viewed as the Ising Network Opinion Formation (INOF) model. In fact this INOF model is directly suitable for the modeling of fibrosis progression in the PPI MetaCore network and we are using it here with certain slight modifications described in the next Section. We call this modified system as the Ising Network FIbrosis (INFI) model. It should be pointed out that a somewhat similar asynchronous Monte Carlo process is used in models of associative memory (see e.g. [27,28]).

In this work, using the results reported in [5,12], we take in the INFI model an initial state of 10 fixed red nodes that produce positive activation response of fibrosis on growth factor-*β* (TGF-*β*) considering them as fixed 10 red protein nodes. We also consider 6 (or 10, 14) fixed blue protein nodes with negative response (see details in next Section). As a result of fibrosis progression via asynchronous Monte Carlo iterations we obtain the steady-state of affected and healthy protein nodes with about 60 - 80 percents of proteins affected by fibrosis. This corresponds to a strong fibrosis activation resolution of a large part of the system.

To reduce this strong fibrosis activation we develop an efficient barrage approach based on the analysis of Erdös proteins linked with fixed red nodes. We show that this Erdös barrage constructed in a clever manner allows to obtain a striking reduction of fibrosis activated resolution nodes by a factor of 300 so that almost all protein nodes are transformed in the healthy phase. We hope that the barrage strategy developed in this work will find applications in clinical experiments with fibrosis progression. The paper constructing as follows: Section 2 describes the data sets, construction of Markov chain transitions, Google matrix and the INFI model, Section 3 presents the obtained results and Discussion and conclusion are given in Section 4. In the Appendix we provide additional data and Figures.

## 2. Data sets and INFI model description

### 2.1 Network data sets

In this work we use the same global PPI MetaCore network as in [12]. It contains *N* = 40079 nodes with *N*_*𝓁*_ = 292191 links (without self connections which existed in [11]). The number of directed activation/inhibition links is *N*_*𝓁*+_/*N*_*𝓁*−_ = 65157/49321 ≃ 1.3 and the number of neutral directed links is *N*_*𝓁n*_ = *N* − *N*_*𝓁*+_ − *N*_*𝓁*−_ = 177713. Here we do not take into account the bi-functional activation/inhibition nature of links. Thus we simply have a directed network with *N* nodes and number of directed links being *N*_*𝓁*_.

For convenience, we present in Table 1, taken from [12], 54 selected fibrosis proteins (nodes). These nodes are composed with 4 TGF-*β* proteins/nodes (*K*_*t*_ = 1, 2, 3, 4), 20 “up-proteins” (*K*_*u*_ = 1, …, 20), 20 “down-proteins” (*K*_*d*_ = 1, …, 20), both obtained from experiments [5] (as described above) and 10 new “X-proteins” (or “X-nodes”; *K*_*x*_ = 1, …, 10) which have a significant influence on the other 44 nodes. The 4 TGF-*β* nodes correspond to different isoforms of this protein. As in [12] we have 4 groups of proteins and we use a specific index for each group: TGF-*β* proteins with index *K*_*t*_ = 1, 2, 3, 4; up-proteins with a strongest positive response noted by index *K*_*u*_ = 1, · · ·, 20 (ordered by the positive response with the strongest response for *K*_*u*_ = 1); down-proteins with a strongest negative response noted by index *K*_*d*_ = 1, · · ·, 20 (ordered by the modulus of negative response with the strongest response modulus for *K*_*d*_ = 1); external proteins noted by index *K*_*x*_ ordered by their local PageRank index (strongest PageRank probability of these 10 proteins is at *K*_*x*_ = 1; see more detail below). All these 54 proteins have their global index *K*_*g*_ = 1, · · ·, 54 used in Table 1.

**Table 1.** Table of the subset of *N*_*r*_ = 54 selected fibrosis proteins (nodes). Here *K*_*g*_ represents the global index of this group, *K*_*t,u,d,x*_ represent the index of the four subgroups of 4 TGF-*β* proteins, 20 up-proteins, 20 down-proteins and 10 additional X-proteins; the *K* (*K*^∗^) index represents the PageRank (CheiRank) index for the global MetaCore network of *N* = 40079 nodes; the last column gives the associated protein names.

| $K_g$ | $K_{t,u,d,x}$ | $K$ | $K^*$ | Protein |
| --- | --- | --- | --- | --- |
| 1 | $K_t = 1$ | 10780 | 26299 | TGF-β 0 |
| 2 | $K_t = 2$ | 235 | 5690 | TGF-β 1 |
| 3 | $K_t = 3$ | 968 | 25073 | TGF-β 2 |
| 4 | $K_t = 4$ | 4726 | 29508 | TGF-β 3 |
| 5 | $K_u = 1$ | 28737 | 25928 | ADAMTS16 |
| 6 | $K_u = 2$ | 3478 | 25137 | FGF21 |
| 7 | $K_u = 3$ | 40048 | 28152 | TNFSF18 |
| 8 | $K_u = 4$ | 2467 | 19160 | ACAN |
| 9 | $K_u = 5$ | 1489 | 24511 | RPH3A |
| 10 | $K_u = 6$ | 26600 | 29559 | ADAMTS8 |
| 11 | $K_u = 7$ | 34769 | 39960 | MEGF6 |
| 12 | $K_u = 8$ | 26295 | 27326 | SV2B |
| 13 | $K_u = 9$ | 27111 | 36021 | C1QTNF3 |
| 14 | $K_u = 10$ | 34616 | 39841 | ANO4 |
| 15 | $K_u = 11$ | 12696 | 16566 | IL11 |
| 16 | $K_u = 12$ | 26624 | 23640 | CDH10 |
| 17 | $K_u = 13$ | 7263 | 30243 | HTR2B |
| 18 | $K_u = 14$ | 4647 | 6551 | LAMA1-1 |
| 19 | $K_u = 15$ | 8342 | 26295 | LAMA1-2 |
| 20 | $K_u = 16$ | 4021 | 8252 | RAPGEF4 |
| 21 | $K_u = 17$ | 29945 | 36964 | DNER |
| 22 | $K_u = 18$ | 22159 | 8569 | GALNT3 |
| 23 | $K_u = 19$ | 29145 | 15531 | ACSBG1 |
| 24 | $K_u = 20$ | 24786 | 8735 | OLFM2 |
| 25 | $K_d = 1$ | 19039 | 28262 | CLEC3B |
| 26 | $K_d = 2$ | 26477 | 28290 | SCARA5 |
| 27 | $K_d = 3$ | 26109 | 11185 | SLC10A6 |
| 28 | $K_d = 4$ | 6360 | 29204 | CXCL5 |
| 29 | $K_d = 5$ | 14952 | 8729 | MYOC |
| 30 | $K_d = 6$ | 5961 | 22288 | IFITM1 |
| 31 | $K_d = 7$ | 5599 | 4483 | ANGPTL4 |
| 32 | $K_d = 8$ | 25538 | 17434 | SELENBP1 |
| 33 | $K_d = 9$ | 18938 | 33179 | FMO1 |
| 34 | $K_d = 10$ | 34080 | 39427 | GPR88 |
| 35 | $K_d = 11$ | 6276 | 22141 | HMGCS2 |
| 36 | $K_d = 12$ | 37060 | 28328 | LGI2 |
| 37 | $K_d = 13$ | 9162 | 2485 | PTN |
| 38 | $K_d = 14$ | 513 | 5974 | ADORA2A |
| 39 | $K_d = 15$ | 7789 | 22652 | GFRA1 |
| 40 | $K_d = 16$ | 6718 | 8844 | IL1R2-1 |
| 41 | $K_d = 17$ | 35446 | 28306 | IL1R2-2 |
| 42 | $K_d = 18$ | 12148 | 3444 | PEG10 |
| 43 | $K_d = 19$ | 27829 | 36195 | FMO2 |
| 44 | $K_d = 20$ | 1973 | 24994 | COX4I2 |
| 45 | $K_x = 1$ | 3 | 13 | β-catenin |
| 46 | $K_x = 2$ | 4 | 6 | p53 |
| 47 | $K_x = 3$ | 11 | 10 | ESR1 |
| 48 | $K_x = 4$ | 13 | 25 | STAT3 |
| 49 | $K_x = 5$ | 22 | 11 | RelA |
| 50 | $K_x = 6$ | 38 | 82 | PPAR-γ |
| 51 | $K_x = 7$ | 111 | 767 | IKK-β |
| 52 | $K_x = 8$ | 179 | 198 | SNAIL1 |
| 53 | $K_x = 9$ | 237 | 1520 | MMP-14 |
| 54 | $K_x = 10$ | 578 | 2123 | Flotillin-1 |

### 2.2 Without formulas: methods, characteristics and expected network results

As in [12] here we present qualitative explanations without formulas of the mathematical methods and characteristics described in the next Subsections. Our aim here is to give a global view on our approach for a common reader.

We use the MetaCore directed network [8] which represents an action of a protein A on a protein B in a form of a directed link (edge) for *N* = 40079 proteins forming the network nodes (proteins). These links are obtained on the basis of careful and detailed analysis of scientific literature about thousands of experiments of various research groups that allowed to collect information about PPI and thus generated a network database with *N* = 40079 nodes and *N*_*𝓁*_ = 292191 links. Certain medical applications of the MetaCore network can be found in [9,10].

The universal mathematical methods to analyze such networks are generic and based on the concept of Markov chains and Google matrix [13–15]. The validity of these methods has been confirmed for various directed networks from various fields of science. Therefore, since the Google matrix analysis is based on a generic mathematical foundation, we expect that this analysis will also work efficiently for PPI networks.

The Google matrix of the global MetaCore PPI network *G* is constructed with specific rules briefly described in the next Subsection 2.3 and in detail in [13–15]. The matrix *G* is obtained from a matrix of Markov chain transition elements *S*_*ij*_ that give weights of transitions between nodes. The important property of *G* is that its application (multiplication) to an arbitrary initial vector *v* preserves the probability and the normalization of this vector (sum of all vector elements) remains constant (taken to be unity). As a result of multiple multiplications by *G* any initial vector converges to the steady-state distribution given by the PageRank vector *P*. The components of this vector represent the probabilities of each node (protein) in this limit. The nodes with the highest probabilities are the most influential nodes of the network (all nodes are monotonically ordered by decreasing values of the PageRank components which provides the “PageRank index” *K* such *K*(*j*) = 1, 2, … for nodes *j* with largest values *P*(*j*)). These nodes have typically many ingoing links and it is likely that some of these ingoing links come from other nodes that also have large PageRank values.

For the inverse network, in which all link directions are inverted, the corresponding PageRank is called CheiRank vector *P*^∗^ [15]. The highest probabilities *P*^∗^ (*j*) are for nodes *j* with the CheiRank index *K*^∗^ (*j*) = 1, 2, … being the most communicative nodes. They typically have many outgoing links. In the INFI model Ising spins with polarization *±*1 (red or blue) or 0 (white) are placed at each node of network. They describe fibrosis activated protein state (red or *σ*_*j*_ = +1) which can activate other proteins, healthy protein state (blue or *σ*_*j*_ = −1) which can prepare other proteins and neutral state proteins (white or *σ*_*j*_ = 0) that cannot affect other proteins but can be activated or repaired by red or blue proteins. The initial spin configuration has certain fixed red proteins with index *K*_*g*_ = 1, …, 10 (see Table 1) and fixed blue proteins with index *K*_*g*_ = 25, …, 30 (or 34 or 38) whose colors always remain unchanged, all other proteins are white (or *σ*_*j*_ = 0). These white proteins, and also red or blue non-fixed proteins, can change their color depending on the majority color of nodes directly linked to it (taking into account the weights of ingoing and outgoing links of the matrix *S*_*ij*_ taken without dangling nodes). This choice is similar to social relations when a member of society takes his opinion to be imposed by opinions of other members related with him. An asynchronous Monte Carlo process performs multiple changes of spins leading to the convergence to a steady-state polarization of all spins. The fraction of red spins *f*_*r*_ determines if the fibrosis progression affected a major part of network or not. We give the mathematical definitions of the INFI model in next Subsections.

### 2.3 Markov chains, Google matrix, PageRank and CheiRank

In this section we remind some basic mathematical definitions of Markov chains and how the Google matrix and the related PageRank and CheiRank vectors for the MetaCore network are constructed. For this we use a simple version of the MetaCore network where initial links are not weighted (see Refs. [11,12] for some details about a more refined version with bi-functional links).

First, one introduces the adjacency matrix with elements *A*_*ij*_ = 1 if a node *j* points to another node *i* (different from *j*) and *A*_*ij*_ = 0 if there is no link from *j* to *i* or if *i* = *j* (we use the same convention as in Ref. [12] where all diagonal elements are chosen to be *A*_*jj*_ = 0 even if there is a link from a node *j* to itself). Then we define a matrix 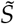 by 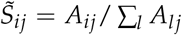 and 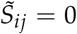 otherwise (i.e. for columns *j* where all elements *A*_*lj*_ are 0). This matrix does not yet describe a Markov process since it has potentially some zero columns *j* (such nodes *j* are also called dangling nodes). However, in this work we will use a Monte Carlo process based on this particular matrix (see next section).

To obtain a proper stochastic matrix *S* one replaces *S*_*ij*_ = 1/*N* for the dangling nodes *j* and 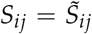 for the other nodes (dangling nodes have no outgoing links so that a column of dangling node *j* has only zero elements in the matrix 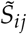). Here *S*_*ij*_ represents the transition probability for a random surfer from node *j* to *i* and the column sum normalization ∑_*i*_ *S*_*ij*_ = 1 ensures the conservation of probability.

The Google matrix elements *G*_*ij*_ are typically defined by

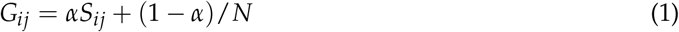

where *α* = 0.85 is the usual damping factor [13,14]. The Google matrix is also a proper stochastic matrix (column sum normalized) and here the random surfer jumps with probability *α* on the network according to *S* and with a probability (1 − *α*) to an arbitrary random node of the network. The damping factor modification helps to avoid possible isolated communities and ensures that the Markov process converges for long times rather quickly to a uniform stationary probability distribution. The latter defines the PageRank vector *P* which is actually the right eigenvector of the Google matrix *G* corresponding to the leading eigenvalue *λ* = 1, i.e. *GP* = *P*. The elements *P*(*j*) of the PageRank vector correspond to the probability to find the random surfer on the node *j* in the stationary limit of the Markov process. The PageRank index *K*(*j*) is obtained by ordering the nodes with decreasing values of *P*(*j*), i.e. the highest (lowest) PageRank probability *P*(*j*) corresponds to *K*(*j*) = 1 (*K*(*j*) = *N*). In this work we will use the *K*-rank of a node to characterize this node in a unique way. The PageRank probability *P*(*j*) is typically related to the number of ingoing links pointing to node *j*. However, it also takes into account the “importance” (i.e. PageRank probability) of the nodes having a direct link to *j*. One can also consider the network obtained by the inversion of all link directions. The same construction as described above provides a Google matrix noted as *G*^∗^ and the corresponding PageRank vector is called the CheiRank vector *P*^∗^, defined by *G*^∗^*P*^∗^ = *P*^∗^. The CheiRank probability *P*^∗^ (*j*) is typically related to the number of outgoing links weighted by the value *P*^∗^ (*i*) of nodes *i* having a link from *j* to *i*. The CheiRank index *K*^∗^ (*j*) is obtained by ordering the CheiRank vector with decreasing values of *P*^∗^ (*j*).

For further details about the properties of Google matrix, PageRank and CheiRank vectors, with many different example networks (also with proteins networks), we refer to [11,12,14,15].

### 2.4 Ising spin network, Monte Carlo process for INFI model

In this work, we consider an Ising type of model using the symmetrized matrix 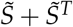 to model the effective spin interactions. We denote by *σ*_*i*_ the spin associated to the node (protein) *i* but here we allow for three different values being *σ*_*i*_ = +1 (“red state”, “fibrosis state”), *σ*_*i*_ = −1 (“blue state”, “healthy state”) and also *σ* = 0 (“white state”, “undetermined or neutral state”). We study this model using a specific Monte Carlo process similar to [26]. The white state will only be used in the initial condition and is supposed to be only temporary.

First, we fix 10 specific nodes permanently to the red state, those with *K*_*g*_ = 1, …, *K*_*g*_ = 10 in the set of Table 1 corresponding to the four TGF proteins and the first 6 *K*_*u*_-proteins with strongest positive response on the TGF stimulation [12]. We also fix *n*_*b*_ nodes permanently to the blue state being the nodes with *K*_*g*_ = 25, …, *K*_*g*_ = 24 + *n*_*b*_ corresponding to the first *n*_*b*_ *K*_*d*_-proteins with strongest negative TGF-response. Here we choose mostly *n*_*b*_ = 6 (same number as the 6 *K*_*u*_ proteins with fixed red state) but we also present some data for the cases *n*_*g*_ = 10 (same total number of fixed red proteins) and *n*_*g*_ = 14. Therefore 10 + *n*_*b*_ nodes have already a permanently fixed spin value *σ*_*i*_ = *±*1.

We implement a Monte Carlo procedure that modifies the other *N*_*v*_ = *N* − 10 − *n*_*b*_ remaining nodes with “variable” spin values (non fixed). For this at given iteration time *τ*, we update the *N*_*v*_ variable nodes by computing the sum over all nodes *j* of the network (fixed and variable) :

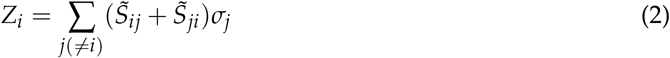

and choosing *σ*_*i*_(*τ* + 1) = +1 if *Z*_*i*_ *>* 0, *σ*_*i*_(*τ* + 1) = −1 if *Z*_*i*_ *<* 0 and *σ*_*i*_(*τ* + 1) = *σ*_*i*_(*τ*) (unchanged) if *Z*_*i*_ = 0. Without going into technical details, we mention that the numerical implementation of (2) exploits of course the sparse structure of the matrix 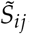. This update is done for every node *i* in the set of *N*_*v*_ variable nodes in a random order using a random permutation (thus no repetitions) which is chosen at the beginning of the iteration process. Once a spin value for a node *i* has been updated, the new value *σ*_*i*_(*τ* + 1) will be used for the update of the subsequent nodes 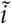. We note that somewhat similar asynchronous Monte Carlo process what used for bio-networks in [29,30] but there the matrix elements 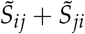 had values *±*1 and network sizes were relatively small (about 100).

This full update procedure for one time step and the full set of *N*_*v*_ variable nodes is repeated up to the iteration time *τ*_max_ = 100 (with different random permutations for the update order at each time step). We also verify if at any given value of *τ* there is ideal convergence where the spins no longer change. (This happens typically at *τ*_*last*_ ≈ 10; see below for details).

The full iteration procedure with *τ*_max_ = 100 time steps is repeated *R* = 100000 times with different random permutations and eventually also different random positions of initial blue or white nodes (depending on the precise type of initial condition).

During this procedure, we compute the fraction of red outcome *f*_*r*_(*i*) for each node as the number of times the node *i* has red state at *τ*_max_ (at different realizations) divided over *R* and also the overall average *f*_*r*_ = ∑_*i*_ *f*_*r*_(*i*)/*N*_*v*_ over the nearly full network of variable nodes. To test the convergence, we also compute the overall average at intermediate values of *τ*.

In order to test the procedure and also in the direct influence of the permanently fixed nodes, we first choose an initial condition where all *N*_*v*_ variable nodes are initialized to the white state. Figure A1 illustrates for this case the convergence of (the overall network average) *f*_*r*_(*τ*) with iteration time *τ*. The convergence seems to be very good at *τ >* 5 with nearly constant values on graphical precision. However, a closer look at the data shows that | *f*_*r*_(*τ* + 1) − *f*_*r*_(*τ*)| ∼ 10^−5^ for *τ* = 10 and the value of *f*_*r*_(*τ*) becomes constant only at *τ* ≥ *τ*_*last*_ ≈ 70 with typical last non-vanishing differences | *f*_*r*_(*τ*_*last*_ − 1) − *f* (*τ*_*last*_)| ∼ 10^−10^ and | *f*_*r*_(*τ*_*last*_ + 1) − *f* (*τ*_*last*_)| = 0. We have verified that this behavior is due to the fact that typically at *τ* ≈ 10 there is exact convergence for a given realization of the random permutations (random pathways) with no further modifications of the spins at *τ >* 10. However, there are rare realizations with larger values, up to *τ* = *τ*_*last*_ ≈ 70 for exact convergence.

In the next section, we present the results for different quantities and also other initial conditions.

## 3. Results

### 3.1 Numerical results

All results presented in this work were obtained with maximal iteration time *τ*_*max*_ = 100 and *R* = 100000 realizations for (potential) different initial conditions and random permutations for the Monte Carlo procedure. Also when we describe a particular initial condition it is implicitly understood that the permanently fixed 10 red and *n*_*b*_ blue nodes given above are indeed fixed to these values from the very beginning, i.e. if we say “that there are *n*_*ib*_ initial blue values” (with *n*_*ib*_ = 0, 1, …) we mean that there are *n*_*ib*_ blue nodes *on the set of N*_*v*_ *variable (non-fixed) nodes*, other variable nodes are initialized to white values and the fixed nodes have still their 10 red and *n*_*b*_ blue values (in particular the value of *n*_*ib*_ does not include the number of *n*_*b*_ fixed blue nodes).

In Figure 1, we show for the case of white or zero initial spins (for the *N*_*v*_ variable nodes; i.e. *n*_*ib*_ = 0) and the three cases of *n*_*b*_ = 6, 10, 14 initial blue fixed nodes (and 10 fixed nodes as explained above) the probability distribution (normalized by an integral) of fraction of red outcomes *f*_*r*_(*i*) for the nodes obtained by a histogram with bin width Δ *f*_*r*_ = 0.01. These distributions are strongly peaked at values close to *f*_*r,peak*_ ≈ 0.93 (*n*_*b*_ = 6), *f*_*r,peak*_ ≈ 0.81 (*n*_*b*_ = 10), *f*_*r,peak*_ ≈ 0.65 (*n*_*b*_ = 14) for roughly 82 % of nodes. The secondary peak at *f*_*r*_ = 0 corresponds to the fraction (number) ≈ 0.18 (7230) of nodes (same value for the three cases of *n*_*b*_) which stay white after 100 iterations for all *R* = 100000 pathway realizations. This set of stable white nodes correspond to nodes not connected to the small number of fixed red and blue nodes. However, despite the small value of only 10 initial fixed red notes the majority of the other nodes has a red outcome, especially for *n*_*b*_ = 6 and with slightly reduced values for *n*_*b*_ = 10, 14. Note that the bin width Δ *f*_*r*_ = 0.01 in Figure 1 is still the rather large and histogram computations with Δ *f*_*r*_ = 0.001 and Δ *f*_*r*_ = 0.0001 show that the actual peaks of the distributions are much sharper as visible in Figure 1.

**Figure 1.**
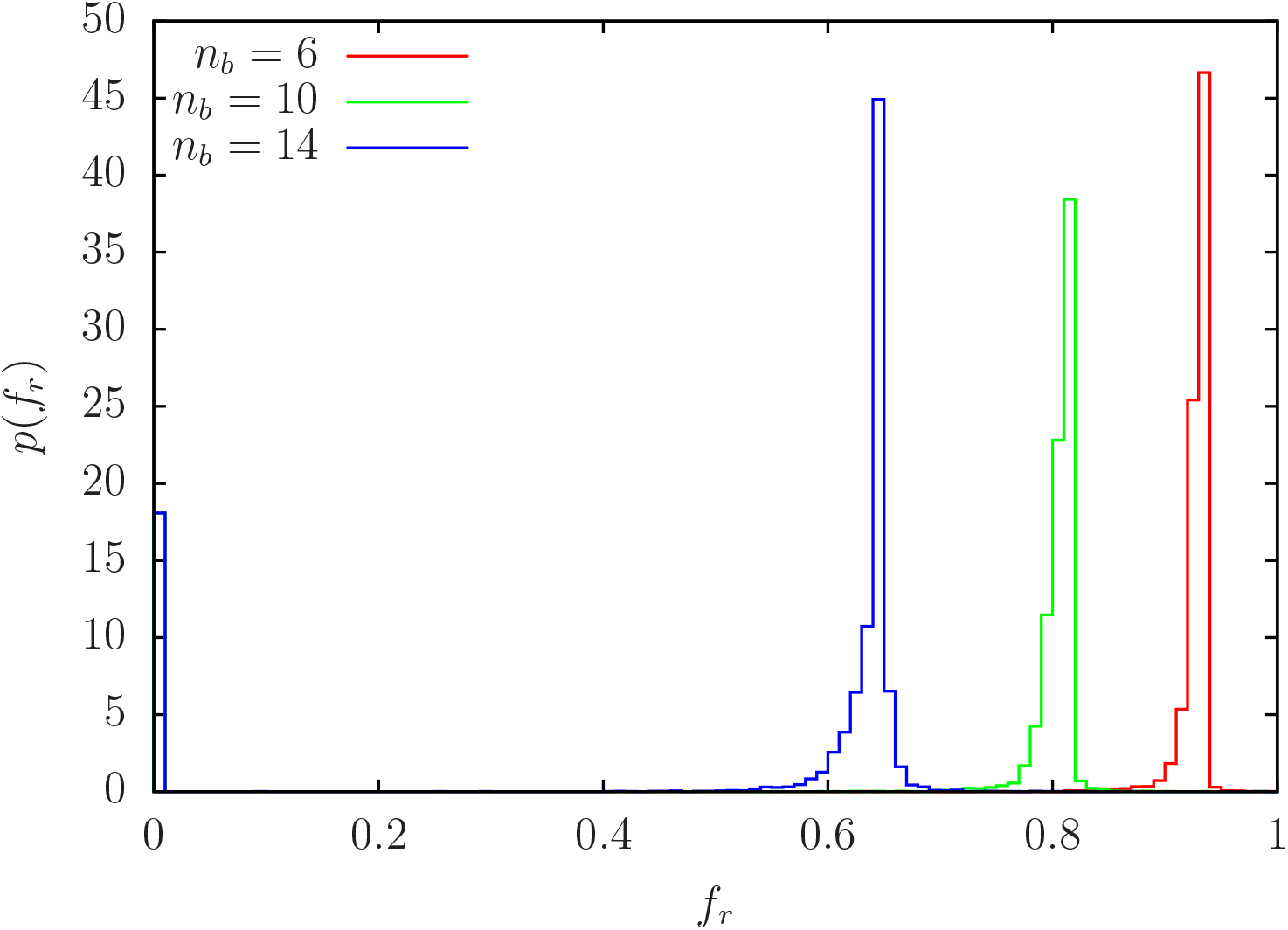
Probability density *p*(*f*_*r*_) of *f*_*r*_ for *n*_*b*_ = 6, 10, 14 obtained from a histogram with bin width Δ *f*_*r*_ = 0.01 using *N*_*v*_ = *N* − 10 − *n*_*b*_ data points *f*_*r*_ (*i*) corresponding to the number of variable nodes. For each node *i* the value *f*_*r*_ (*i*) is obtained as the fraction of red outcome (*σ*_*i*_ = +1) of *R* = 100000 pathway realizations for this node using the initial condition with no initial (variable) blue nodes, i.e. *n*_*ib*_ = 0. The distributions are normalized by 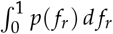 and there are strongly peaked at values close to *f*_*r,peak*_ ≈ 0.93 (*n*_*b*_ = 6), *f*_*r,peak*_ ≈ 0.81 (*n*_*b*_ = 10), *f*_*r,peak*_ ≈ 0.65 (*n*_*b*_ = 14) for roughly 82 % of nodes. The secondary peak at *f*_*r*_ = 0 corresponds to the fraction ≈ 0.180 of nodes (same value for the three values of *n*_*b*_) which stay white after 100 iterations for all *R* = 100000 pathway realizations.

It seems that with the white initial condition (except for the fixed nodes) nearly all nodes (except those in the stable white set) have a large probability for red outcome. Therefore, we try to reduce this red outcome by choosing a certain number *n*_*ib*_ of initial blue (variable) nodes and other nodes with white initial values. The question is also where to place these blue nodes. In a first model we choose the *n*_*ib*_ blue nodes at different random positions (for each of the *R* = 100000 pathway realizations) on the full set of variable nodes giving a set of *N*_*r*_ = *N*_*v*_ of possible blue initial nodes with possible values *n*_*ib*_ = 0, …, *N*_*r*_ and a corresponding fraction *f*_*ib*_ = *n*_*ib*_/*N*_*r*_. Figure 2 shows for the case *n*_*b*_ = 6 (the results for *n*_*b*_ = 10 and *n*_*b*_ = 14 are very similar) the overall network average *f*_*r*_ at maximum iteration time as a function of *f*_*ib*_ (or *n*_*ib*_). The initial value *f*_*r*_ ≈ 0.75855 at *n*_*ib*_ = 0 corresponds to the average which can also be obtained from the histogram data shown in Figure 1 which is roughly 0 *×* 0.18 + 0.93 *×* 0.82 ≈ 0.76. With increasing *f*_*ib*_ (or *n*_*ib*_) the value of *f*_*r*_ decreases, e.g. *f*_*r*_ ≈ 0.1 for *n*_*ib*_ ≈ 60 and *f*_*r*_ ≈ 10^−2^ for *n*_*ib*_ ≈ 100. However, this decrease is not optimal since the initial *n*_*ib*_ blue nodes can be on arbitrary random positions of available variable nodes. Therefore, we also try different network subsets for the potential initial blue nodes.

**Figure 2.**
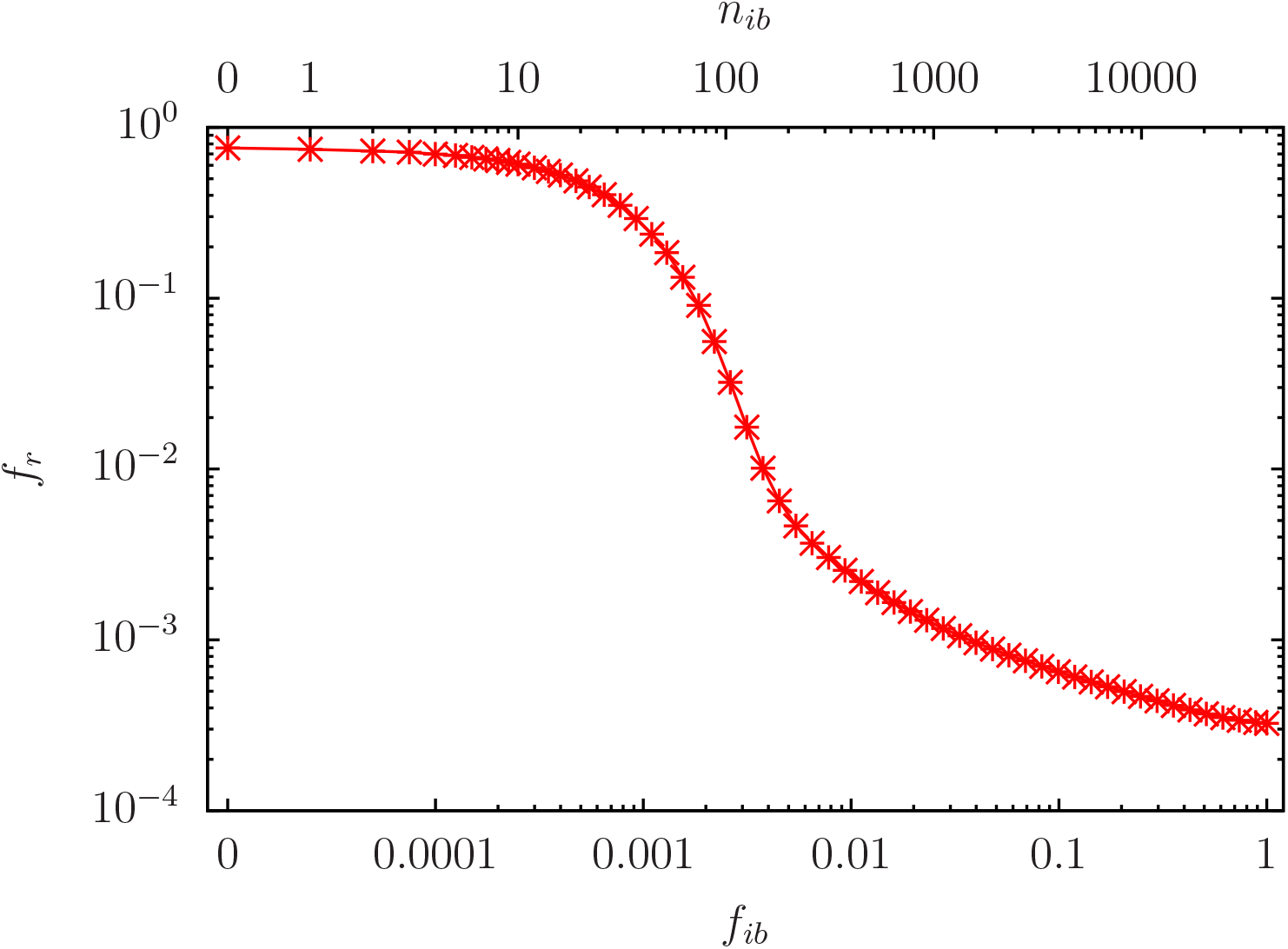
Probability of red outcome *f*_*r*_ = ∑_*i*_ *f*_*r*_ (*i*)/*N*_*v*_ averaged over all (variable) *N*_*v*_ network nodes versus fraction *f*_*ib*_ of random initial blue nodes in the full set of *N*_*r*_ = *N*_*v*_ variables nodes and for the case *n*_*b*_ = 6. The top *x*-axis shows the number *n*_*ib*_ = *N*_*r*_ *f*_*ib*_ of initial (variable) blue nodes. For each value of *n*_*ib*_ the initial condition corresponds to *n*_*ib*_ initial blue nodes (*σ*_*i*_ (*τ* = 0) = −1) with different random positions in the set of size *N*_*r*_ (for the *R* = 100000 pathway realizations). The representation is logarithmic on both axis except for the first data point at *n*_*ib*_ = 0 which has artificially been placed at a finite position below *n*_*ib*_ = 1 for practical reasons. Note that the value *f*_*r*_ (*n*_*ib*_ = 0) ≈ 0.75855 of this data point can also be obtained from the distribution average 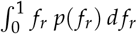 from the data of Figure 1 for the case *n*_*b*_ = 6.

In Figure 3, we show the dependence of *f*_*r*_ on *f*_*ib*_ (or *n*_*ib*_) for a reduced subset of *N*_*r*_ = 38 nodes corresponding to the 38 proteins of Table 1 which are not used for fixed red/blue values, i.e. *K*_*g*_ = 11, …, *K*_*g*_ = 24 and *K*_*g*_ = 31, …, *K*_*g*_ = 54. Now, *n*_*ib*_ represents the number of initial blue nodes on random positions in this subset (and white initial nodes on every other variable node). Now, the decrease of *f*_*r*_ with *n*_*ib*_ seems somewhat stronger, i.e. *f*_*r*_ ≈ 0.1 for *n*_*ib*_ ≈ 8 and *f*_*r*_ ≈ 10^−2^ for *n*_*ib*_ ≈ 16, and we may conclude that this subset is more effective to reduce the red outcome (“to block fibrosis”) than the full set of available variable nodes.

**Figure 3.**
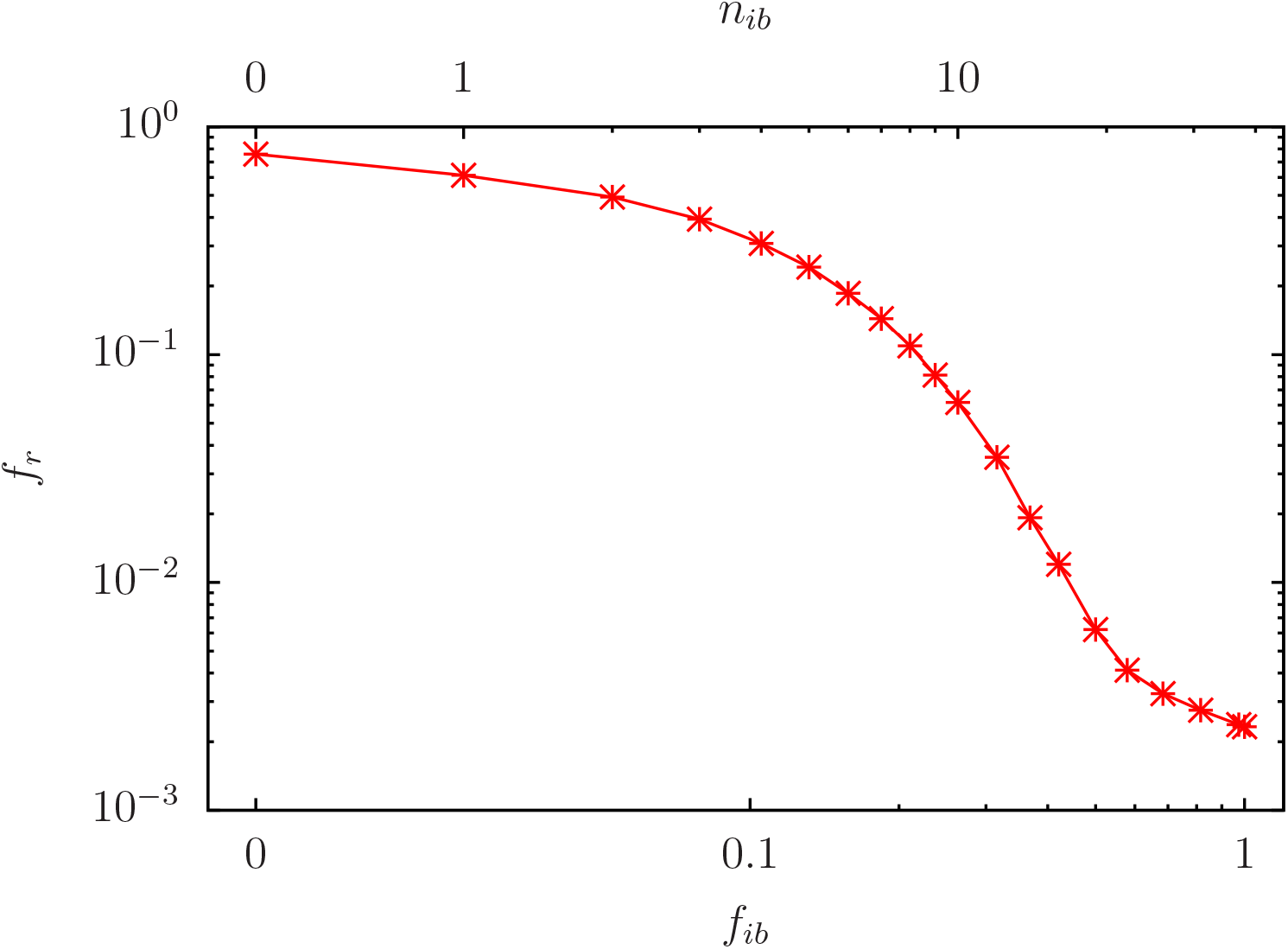
As Figure 2 for *n*_*b*_ = 6 but using a reduced set of potentially initial blue nodes with *N*_*r*_ = 38 using the non-fixed nodes of the set given in Table 1 (i.e. the nodes with *K*_*g*_ = 11, … 24 and *K*_*g*_ = 31, …, 54). For each value of *n*_*ib*_ the initial condition corresponds to *n*_*ib*_ initial blue nodes (*σ*_*i*_ (*τ* = 0) = −1) in this reduced set of 38 nodes (with random positions for each of the *R* = 100000 pathway realizations). All other variable nodes have initial white values (*σ*_*i*_ (*τ* = 0) = 0). The first data point at *n*_*ib*_ = 0 has the same value as in Figure 2 and it has also artificially been placed at a finite position below *n*_*ib*_ = 1 for practical reasons.

To determine a still more effective subset, we determine all nodes which have a direct link or inverse link to one of the 10 fixed red nodes (those with *K*_*g*_ = 1, …, *K*_*g*_ = 10 in Table 1). This defines a particular set, of size *N*_*E*_ = 353, with Erdös number being unity with respect to the 10 fixed red nodes as HUB and using the symmetrized link matrix 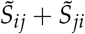. Figure 4 shows the dependence of *f*_*r*_ on *f*_*ib*_ (or *n*_*ib*_) for this Erdös set with *N*_*r*_ = *N*_*E*_. Now the decrease is even more effective with *f*_*r*_ ≈ 0.1 for *n*_*ib*_ ≈ 6 and *f*_*r*_ ≈ 10^−2^ for *n*_*ib*_ ≈ 14,

In Figure 5, we show in a color plot the dependence of *f*_*r*_(*i*) for the 54 nodes *i* belonging to the set of Table 1 on the index *n*_*g*_ which is a monotonic function of *n*_*ib*_ (essentially linear for *n*_*ib*_ ≤ 10 and logarithmic for *n*_*ib*_ *>* 10; see caption of Figure 5 and Appendix Figure A1 for details). The data of Figure 5 correspond to the data of Figure 4 using the Erdös set with *N*_*E*_ = 353 as potential initial blue nodes (i.e. for each value of *n*_*ib*_ ∈ {0, …, 353} we have *n*_*ib*_ initial blue nodes at random positions in this set). In Fig 5, we can of course identify the 10 fixed red nodes (*K*_*g*_ = 1, …, 10) and the *n*_*b*_ = 6 fixed blue nodes (*K*_*g*_ = 25, …, 30) with either *f*_*r*_(*i*) = 1 or *f*_*r*_(*i*) = 0 respectively for all values of *n*_*g*_. The other nodes follow quite closely the decrease of Figure 4 for the global average of *f*_*r*_. However, for certain specific nodes (*K*_*g*_ = 17, 23, 36, 38, 41, 44) the decay of *f*_*r*_ with *n*_*g*_ (or *n*_*ib*_) is less pronounced for *n*_*g*_ *>* 10. Apparently, these nodes are less likely to be blocked by a modest number of initial blue nodes.

**Figure 4.**
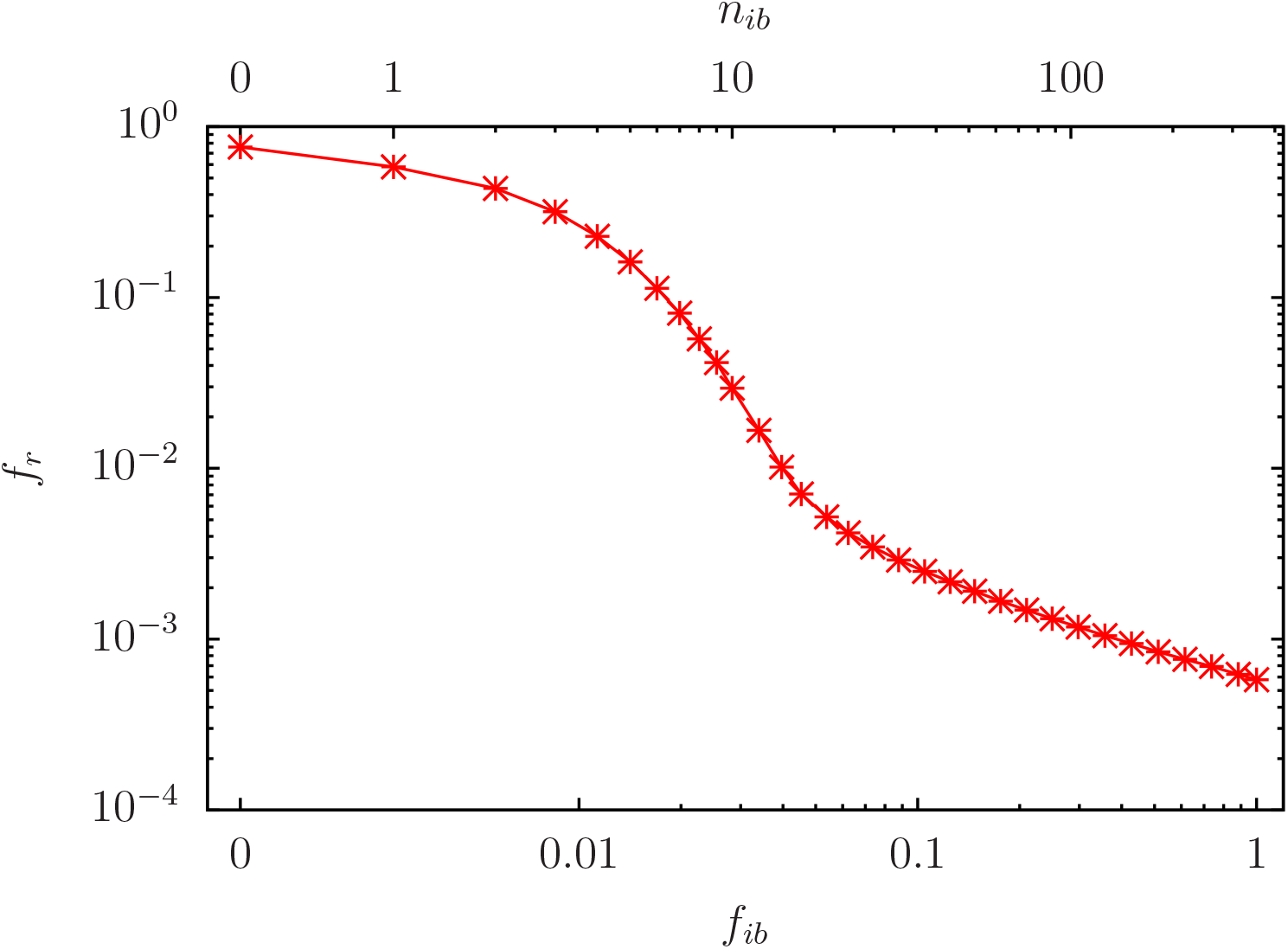
As Figure 3 for *n*_*b*_ = 6 but using the Erdös set (all nodes with direct links in both directions to the 10 fixed red nodes) as reduced set of potentially initial blue nodes with *N*_*r*_ = *N*_*E*_ = 353. The first data point at *n*_*ib*_ = 0 has the same value as in Figure 2 and it has also artificially been placed at a finite position below *n*_*ib*_ = 1 for practical reasons.

**Figure 5.**
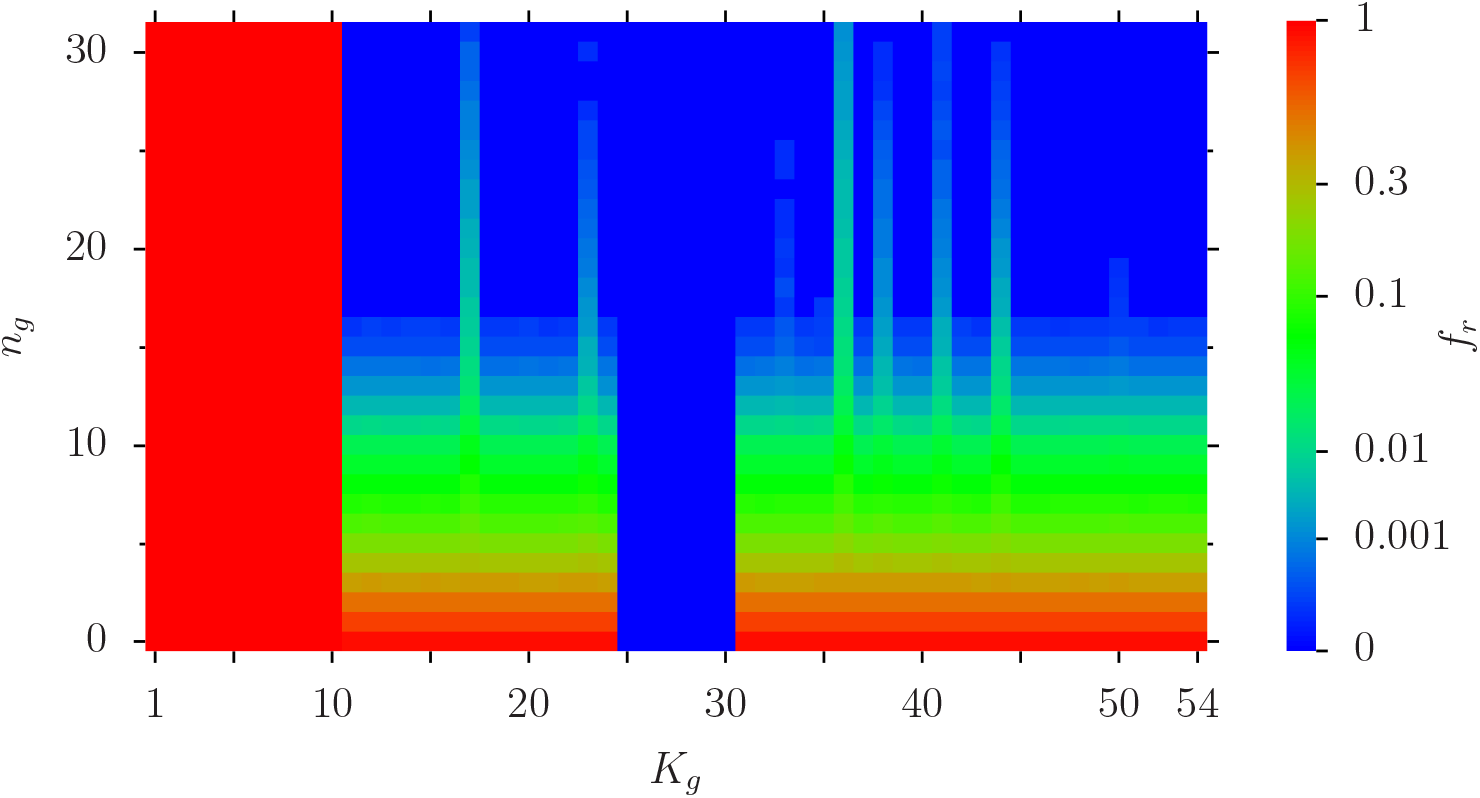
Color plot of *f*_*r*_ (*i*) for the 54 nodes of the set of Table 1 for the case of Figure 4 (i.e. *n*_*b*_ = 6, using the Erdös group with *N*_*r*_ = 353 nodes as potential initial blue nodes and *n*_*ib*_ being the number of initial blue nodes at random positions in the Erdös group). The *x*-axis corresponds to the index *K*_*g*_ of Table 1 and the *y*-axis represents to the coarse-grained index *n*_*g*_ = 0, …, 31 which corresponds to *n*_*ib*_ = *n*_*g*_ for *n*_*g*_ ≤ 10 (linear scale) and 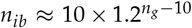 for *n*_*g*_ *>* 10 (logarithmic scale). See also Figure A2 which shows the link between *n*_*ib*_ and *n*_*g*_. The values of the color bar correspond to *f*_*r*_ (*i*) (i.e. red for *f*_*r*_ (*i*) = 1, green for *f*_*r*_ (*i*) ≈ 0.06 and blue for *f*_*r*_ (*i*) = 0). Here small *f*_*r*_ (*i*) values have been amplified to improve the visibility (non-linear scale in the Colombo).

We have also analyzed the data of Figure 2 (using all *N*_*v*_ variable nodes as potential initial blue nodes) and Figure 3 (using the remaining set of 38 non-fixed nodes of Table 1 as potential initial blue nodes) with similar color plots and in both cases we observe the same qualitative behavior as in Figure 5 : identification of fixed red/blue nodes, similar decrease of *f*_*r*_(*i*) with increasing *n*_*ib*_ for the other nodes and less pronounced decrease for the 6 specific nodes mentioned above.

The question arises which of the nodes of the Erdös set, or more generally, which configurations of few selected nodes of this set, are most effective to reduce the red outcome when selected as initial blue nodes. To answer this question, we compute for each node *i* of the Erdös set the red outcome *f*_*rc*_(*i*) averaged over the full network when this specific node *i* is selected as single initial blue node, i.e. with *n*_*ib*_ = 1 but now with different cases of given fixed positions (instead of random positions). Note that this quantity is different from *f*_*r*_(*i*) used in the histogram of Figure 1 which is the probability of red outcome of node *i* (not averaged over the network) with full white initial condition. The nodes of the Erdös set can be ordered with increasing values of the new quantity *f*_*rc*_(*i*) which provides a specific ranking index 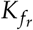 in this set. In Table 2, we present in 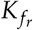 -order the nodes (proteins) of the Erdös set which have either *K* ≤ 40 or *K*^∗^ ≤ 40, either low *K*- or *K*^∗^-rank. It turns out these nodes also provide the lowest values of *f*_*rc*_ and 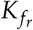 (with some holes 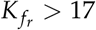).

**Table 2.**
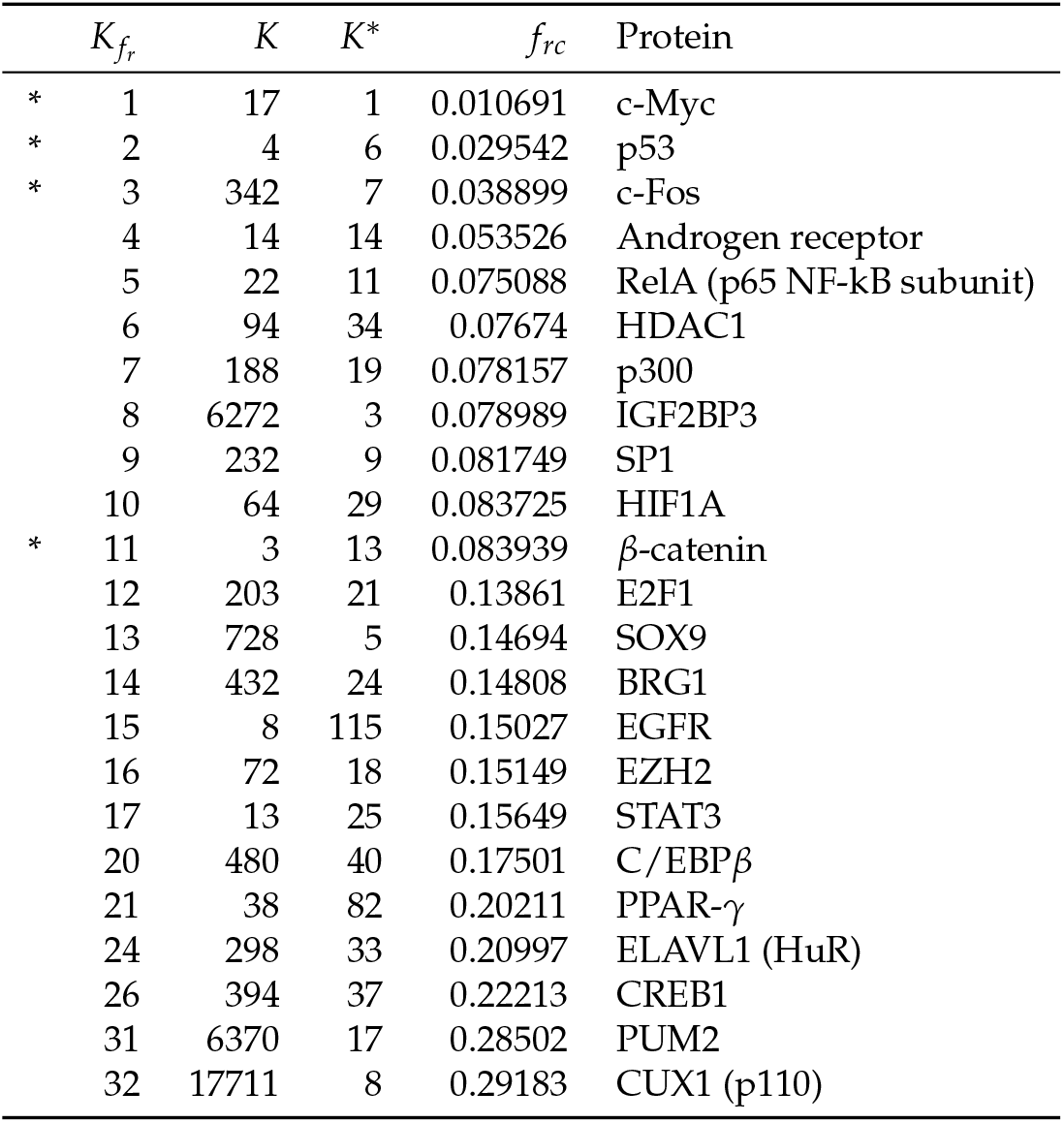
Table of selected fibrosis proteins (nodes) belonging to a smaller subset of the Erdös group such that either *K* ≤ 40 or *K*^∗^ ≤ 40 where the *K* (*K*^∗^) index represents the PageRank (CheiRank) index for the global MetaCore network of *N* = 40079 nodes. *f*_*rc*_ represents the average fraction of red nodes obtained when using the corresponding node as single initial non-fixed blue node and the index 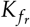 is obtained by ordering the Erdös group of 353 elements with increasing values of *f*_*rc*_; the last column gives the associated protein names. The shown subset of the Erdös group corresponds essentially to the nodes with lowest *f*_*rc*_-value (up to 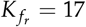).The four nodes marked by “∗” in the first column are the nodes used for the subsequent example computations using 4 specific optimal nodes.

Then we choose as example four “optimal” nodes with *K* = 3, 4, 17, 342 (marked with an asterix in the first column of Table 2). The nodes *K* = 4, 17, 342 occupy indeed the top three places in the 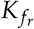-rank while the node *K* = 3 “only” corresponds to the position 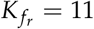. The reason for this choice is related to the fact that these four nodes are more uniformly optimal if we also choose a small number of *n*_*ib*_ = 2, …, 10 initial blue nodes. In this case, using the data of Figure 4 (or the code to produce these data) it is actually possible to compute the conditional probability *f*_*rc*_(*i*) of a red outcome when the node *i* is by chance selected by the random initial condition for *n*_*ib*_ *>* 1 (as one of the initial *n*_*ib*_ blue nodes). For example for *n*_*ib*_ = 4 with *R* = 100000 different random initial conditions of 4 blues nodes out of 353 we have typically a bit more than 1000 realizations where an arbitrary fixed node *i* belongs to the random set of 4 initial nodes. This provides enough data for a reasonable average to compute the conditional probability of red outcome of the node *i*. In this way, it is possible to compute more general 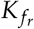-rankings as in Table 2, also for modest values of *n*_*ib*_ *>* 1. It turns out that these rankings produce roughly the same sets of nodes in the first places (with possible permutations between different *n*_*ib*_-values) and the four selected nodes *K* = 3, 4, 17, 342 are indeed optimal as a group for the three values *n*_*ib*_ = 2, 3, 4 (and also some larger values).

In Figure 6, we present (for *n*_*b*_ = 6) the dependence of *f*_*r*_ on *n*_*i*_ for this small optimal set of 4 possible initial blue nodes (red square data points). Here the decrease of *f*_*r*_ with increasing *n*_*i*_ is indeed very strong with *f*_*r*_ ≈ 0.035 for *n*_*ib*_ = 1 and *f*_*r*_ ≈ 0.00318 for *n*_*ib*_ = 4. Furthermore, Figure 6 also provides the conditional probabilities of red outcome 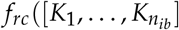 (black small circle data points) for specific configurations 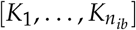 with one configuration for *n*_*ib*_ = 0, 4, four configurations for *n*_*ib*_ = 1, 3 and six configurations for *n*_*ib*_ = 2. We see that there are considerable fluctuations between the different configurations in the red outcome for *n*_*ib*_ = 1 and *n*_*ib*_ = 2 with optimal values being *f*_*rc*_([17]) ≈ 0.009477 and *f*_*rc*_([3, 17]) ≈ 0.004863. Appendix Figure A3 provides two panels of similar figures for the other cases *n*_*b*_ = 10 and *n*_*b*_ = 14.

**Figure 6.**
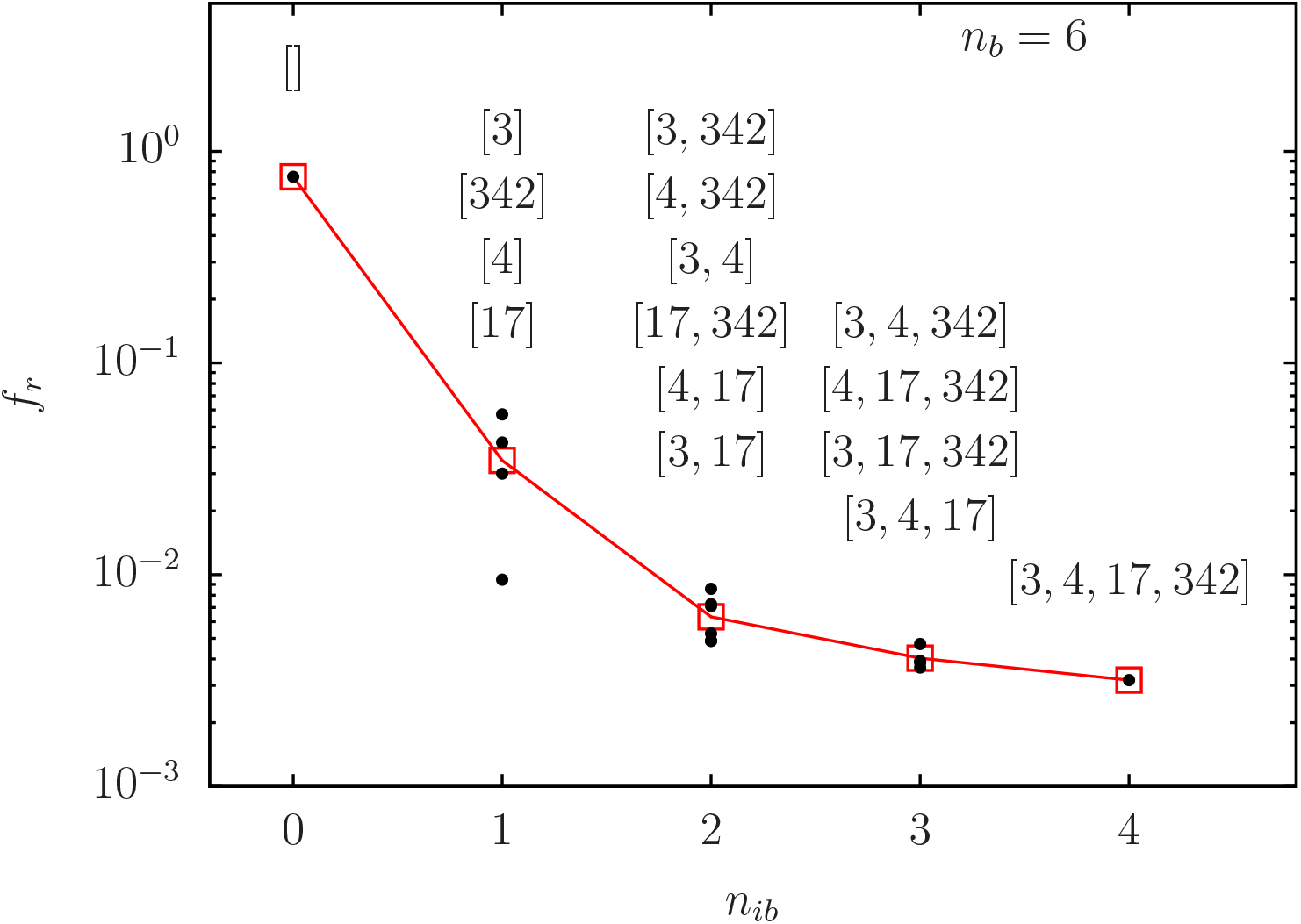
As Figs. 3 and 4 but with linear *x*-axis and using the optimal set with four *K*-values [3, 4, 17, 342] for potential initial blue nodes (i.e. *n*_*b*_ = 6, *N*_*r*_ = 4 and *n*_*ib*_ = 0, …, 4 being the number of initial blue nodes at certain positions in this set). The red square data points represent the average *f*_*r*_ with respect to all possible configurations of *n*_*ib*_ initial blue nodes and the black data points represent the conditional average 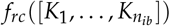 for particular configurations 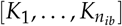 of initial (variable) blue nodes (with *K*_*j*_ being *K*-rank values of nodes and *n*_*ib*_ = 0, …, 4). For each column the top (bottom) black data point corresponds to the top (bottom) configuration shown above. For *n*_*ib*_ = 0 (“empty” configuration “[]” with no initial variable blue node) and *n*_*ib*_ = 4 (full configuration “[3, 4, 17, 342]” of all four nodes) there is only one configuration and therefore only one associated black data point.

Figure 7 is a similar color plot as Figure 5 but for the specific set of 4 optimal nodes [3, 4, 17, 342]. The conclusions are similar to Figure 5 (confirmation of fixed red/blue nodes, strong decrease with increasing *n*_*ib*_ for other nodes and certain specific nodes with less pronounced decrease.

**Figure 7.**
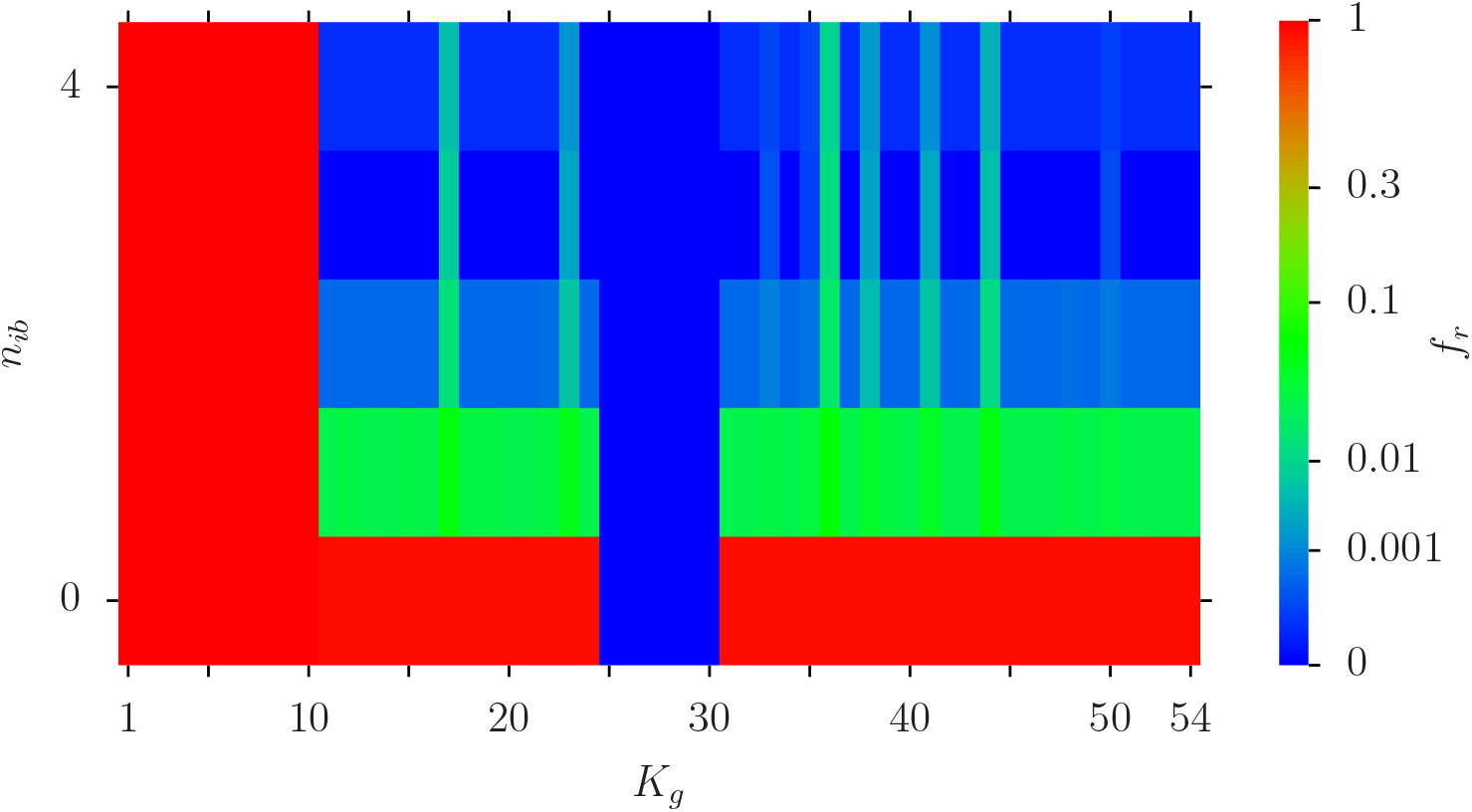
Color plot of *f*_*r*_ (*i*) as in Figure 5 but using the optimal set with four *K*-values [3, 4, 17, 342] for potential initial blue nodes (i.e. *n*_*b*_ = 6, *N*_*r*_ = 4 and *n*_*ib*_ = 0, …, 4 being the number of initial blue nodes at random positions in this set). Here the *y*-axis corresponds directly to *n*_*ib*_ and the *x*-axis to *K*_*g*_ of Table 1. The values of the color bar correspond to *f*_*r*_ (*i*) (with amplified scale as in Figure 5).

In Figure 8, we show the 353 nodes of the Erdös set in the global *K*-*K*^∗^-plane (in a double logarithmic representation) with colored data points such that the color provides the value of *f*_*rc*_(*K*) for each node at given *K*-value (obtained at *n*_*ib*_ = 1 with the corresponding node as initial blue node). Optimal nodes with low red outcome (blue color) have typically small values in their *K*- and *K*^∗^-rank and nodes with large red outcome (red color) have typically large values in their *K*- and *K*^∗^-rank. Furthermore, Figure 8 also provides the positions of the 10 fixed red and the 6 blue nodes in the global *K*-*K*^∗^-plane with typically quite large values for *K* and *K*^∗^. Using the same data Appendix Figure A4 shows the dependence of *f*_*rc*_ (at *n*_*ib*_ = 1) on the rank 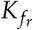. Obviously, this curve is monotonically increasing and the range with small values of *f*_*rc*_ is rather small, i.e. *f*_*rc*_ *<* 0.1 corresponds to 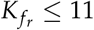 (see also Table 2). Note that the average of the curve in Figure A4 corresponds to the data point at *n*_*ib*_ = 1 in Figure 4 which is *f*_*r*_(*n*_*ib*_ = 1) ≈ 0.5818.

**Figure 8.**
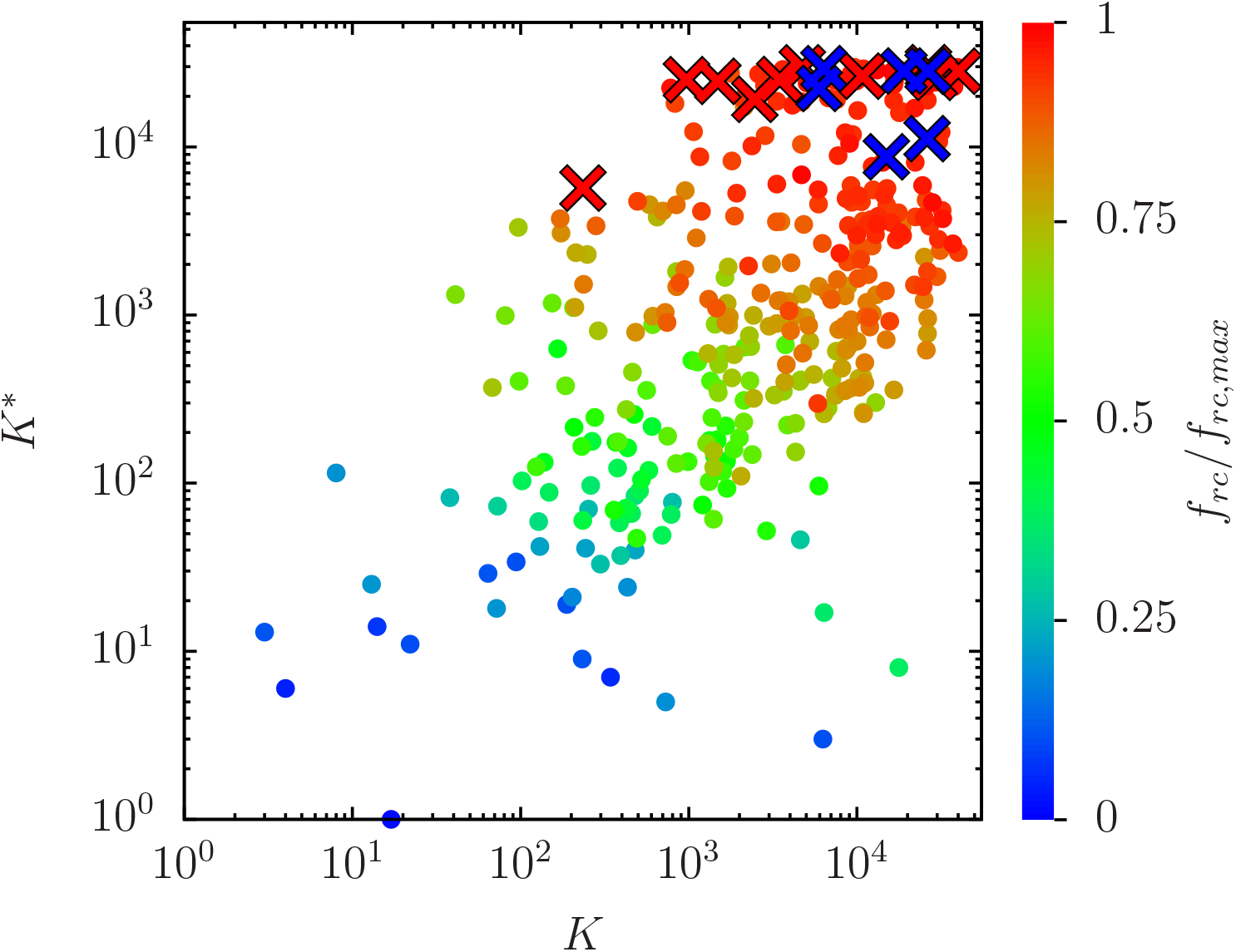
The positions of the 353 nodes of the Erös group in the *K*-*K*^∗^ plane in a double logarithmic representation (colored full circles). The color of each data point corresponds to the value of *f*_*rc*_ (*K*) using the initial configuration [*K*] (i.e. *n*_*ib*_ = 1 with initial blue node at given value of *K*). To be more precise, the values of the color bar correspond to *f*_*rc*_ (*K*)/ *f*_*rc,max*_ with *f*_*rc,max*_ ≈ 0.77775 being the maximum value of *f*_*rc*_ (i.e red for maximum, green for intermediate and blue for zero values and with no amplification of small values). The red (blue) crosses indicate the positions of the fixed 10 red (6 blue) nodes.

We have also analyzed the statistical properties of the individual *f*_*r*_(*i*) values, same quantity as in Figure 4, but using the data of Figure 6 for *n*_*ib*_ = 4 with the initial blue node configuration [3, 4, 17, 342] and global (network averaged) value *f*_*r*_ ≈ 0.00318. For example Appendix Figure A5 shows the 100 nodes with largest values of this quantity in the *K*-*K*^∗^ plane. Obviously, among the 26 nodes with *f*_*r*_(*i*) = 1 we have the 10 permanently fixed red nodes (*K*_*g*_ = 1, …, 10 in Table 1) but there are also further 16 nodes with *f*_*r*_(*i*) = 1 probably with (mostly) exclusive links to the fixed 10 red nodes, i.e. no or few links to other nodes, therefore explaining the fixed outcome *f*_*r*_(*i*) = 1. There are about 1% (4%) of network nodes with values *f*_*r*_(*i*) *>* 0.5 (*f*_*r*_(*i*) *>* 0.08850).

Furthermore, Appendix Figure A6 shows the histogram distribution *p*(*f*_*r*_) for the same data with a rapid decay at *f*_*i*_ *<* 1 and a small peak at *f*_*r*_ = 1 corresponding to the 26 nodes *i* with *f*_*r*_(*i*) = 1. We have also computed the related global probability *P*(*f*_*r*_) for a node *i* to have a value *f*_*r*_(*i*) *> f*_*r*_. This quantity is shown in Appendix Figure A7, confirming the rapid decay which is quite well algebraic as 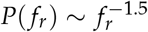 (for *f*_*r*_ ≥ 5 *×* 10^−3^) and which is similar to the Poincare recurrences decay in symplectic chaotic maps with Ulam networks [31].

The slow algebraic decay of *P*(*f*_*r*_) has important consequences. It shows that the average values of *f*_*r*_ shown in Figure 6 even underestimate the effect. Explicitely, the average value of *f*_*r*_ ≈ 0.00318 for *n*_*ib*_ = 4 in Figure 6 corresponds, according to Appendix Figure A7, to a fraction of nodes *P*(0.00318) ≈ 0.22 having an *f*_*r*_(*i*) value above this average and 78% of nodes have a smaller *f*_*r*_(*i*) value. It is well known, that for such long tail distributions one should also focus on the median value *f*_*r,median*_ defined by *P*(*f*_*r,median*_) = 0.5 corresponding in our case to *f*_*r,median*_ = 10^−5^ (only 1 red outcome in the *R* = 100000 pathway realizations) indicating that (slightly more than) 50% of nodes have an *f*_*r*_(*i*) value below or equal to the median value with nearly perfect reduction of the red outcome.

### 3.2 Results without formulas

In this work we presented a mathematical model of fibrosis progression in the PPI MetaCore network describing the global interaction structure of almost all proteins and important molecules (nodes). We show that even with only 10 fibrosis activated proteins the fibrosis progression can spread over a great majority of nodes (about 70 %). The developed analysis of network structure allows us to propose an efficient strategy which allows to reduce a number of fibrosis activated nodes by a factor 300 and disease elimination. The method is based on the Erdös barrage construction: we determine the Erdös nodes directly linked to the fixed 10 activated red nodes; this number can be relatively large (353 in our case); however, we show that a barrage with only 4 blue repairing Erdös nodes, corresponding to the four proteins c-Myc, p53, c-Fos and *β*-catenin, gives a reduction of the average number of fibrosis activated nodes by a factor 300; these 4 nodes belong to network nodes with high PageRank and CheiRank indexes of global MetaCore network.

Furthermore, this average actually underestimates the effect since it is determined by a relative small number of nodes with a modest reduction (e.g. factor of ∼ 100) while for more than 50% of nodes the reduction factor is even 100000 (only 1 infection outcome in the 100000 statistical realizations of our simulation). We have also identified in the group of Table 1 six interesting proteins HTR2B, ACSBG1, LGI2, ADORA2A, IL1R2-2 and COX4I2 (corresponding to the vertical green lines in Figures 5 and 7) where the reduction effect is somewhat less pronounced (compared to the 50% of nodes with nearly perfect reduction). We expect that our INFI model can be tested with other PPI networks (e.g. those of [32,33]).

## 4. Discussion and conclusion

Identifying fibrosis-associated proteins and fibrosis progression is a critical issue in treating heart failure. An experimental determination of the fibrosis progression process is extremely time consuming and labor-intensive. In this work we present and describe a mathematical model of such a fibrosis progression using the global PPI MetaCore network. The performed analysis shows that even a small number of fibrosis activated proteins can lead to a global fibrosis progression of a major part of the whole PPI network.

We developed an efficient method of the Erdös barrage when a small group of e.g. 4 repairing protein can reduce the number of fibrosis activated proteins (nodes) by a factor 300 leading to a healthy state of the global system described here by the MetaCore network. We expect that similar results can be obtained for other disease progression. We think that it would be very interesting to test this INFI approach with other global PPI networks like [32,33]. We hope that the described INFI method will lead to new efficient medical treatments of fibrosis and other various diseases.

## Appendix

### A. Additional figures

Here we present additional Appendix Figures A1,…, A7 for the main part of this article. Appendix Figure A1 illustrates the convergence of the time evolution of spin configurations.

**Appendix Figure A1.**
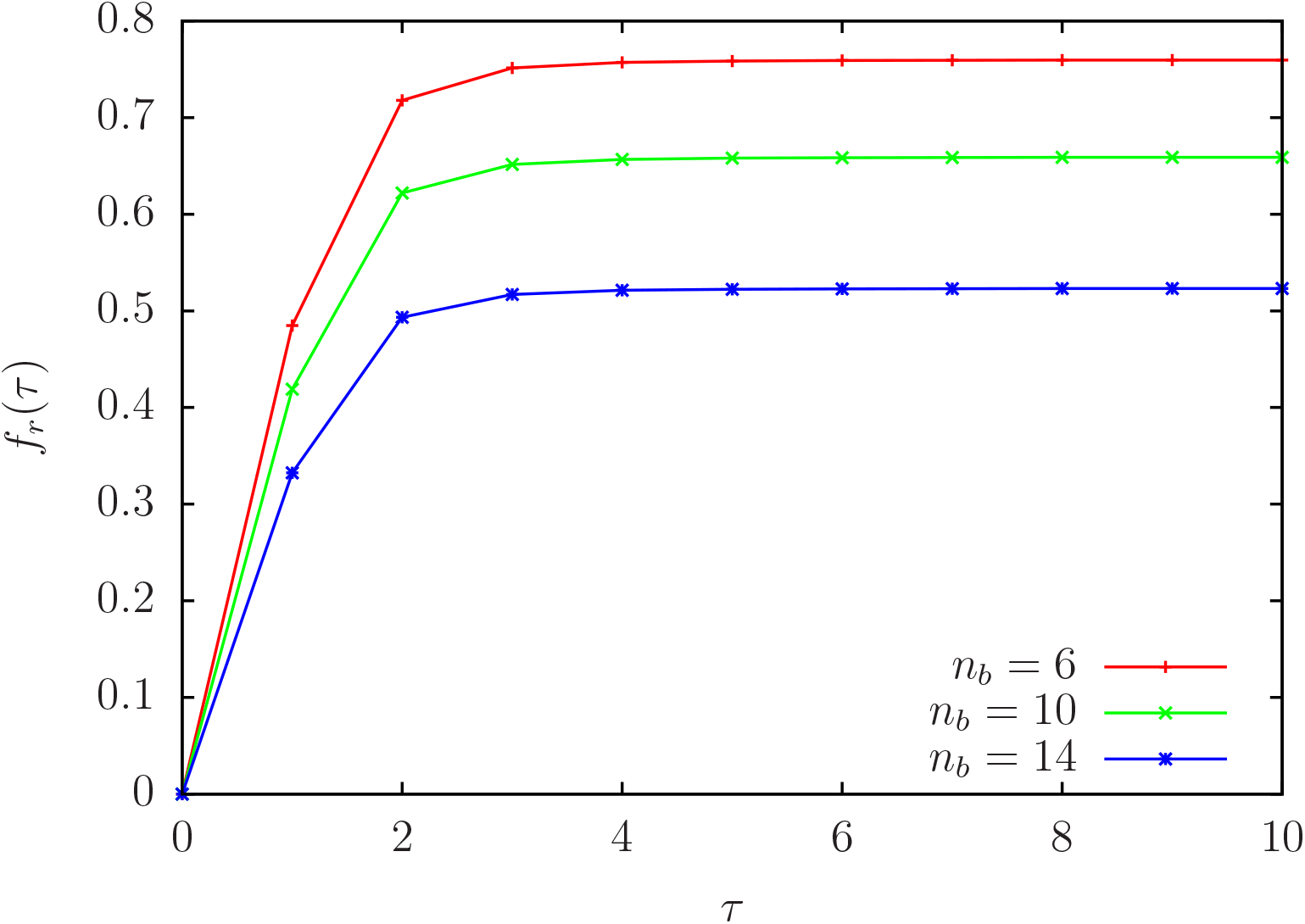
Illustration of the convergence of spin-configurations for *n*_*b*_ = 6, 10, 14 and *n*_*ib*_ = 0. Shown is average *f*_*r*_ (*τ*) versus iteration time *τ*. For practical reasons the convergence seems to be quite good at *τ* ≈ 5. However, the value of *f*_*r*_ (*τ*) becomes constant only at *τ* ≥ *τ*_*last*_ ≈ 70 with typical last non-vanishing differences | *f*_*r*_ (*τ*_*last*_ − 1) − *f* (*τ*_*last*_)| ∼ 10^−10^ and | *f*_*r*_ (*τ*_*last*_ + 1) − *f* (*τ*_*last*_)| = 0. All data in this work have been computed with values up to *τ* = 100 to verify that exact convergence is achieved.

**Appendix Figure A2.**
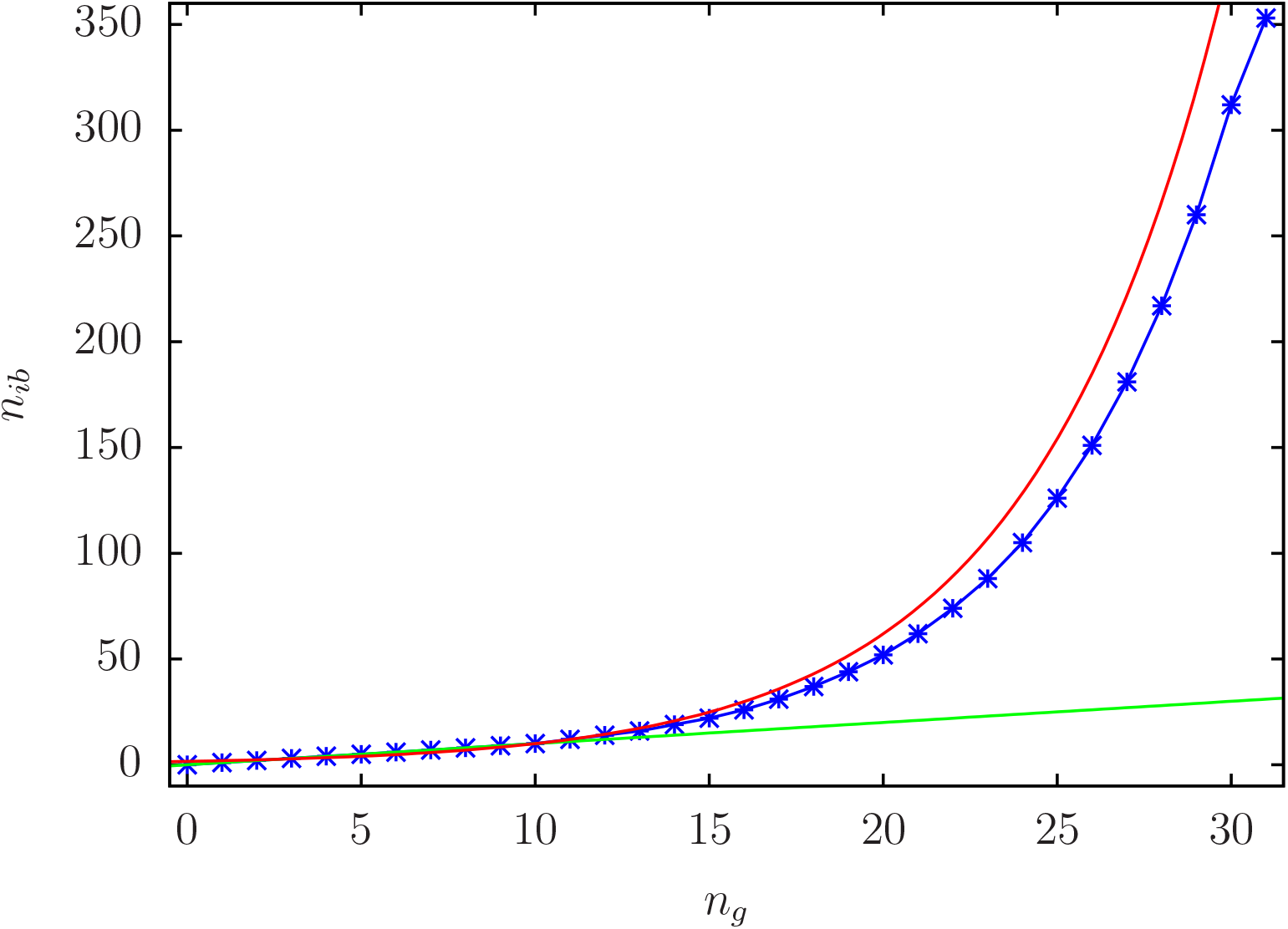
Dependence of *n*_*ib*_ on the coarse-grained index *n*_*g*_ used in Figure 5 (blue/star data points). The green line shows for comparison the linear behavior *n*_*ib*_ = *n*_*g*_ (exactly identical to blue line for *n*_*g*_ ≤ 10) and the red line shows the exponential behavior 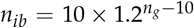 (good approximation for *n*_*g*_ *>* 10).

Appendix Figure A2 illustrates the link between *n*_*ib*_ and the coarse-grained index *n*_*g*_ used in Figure 5.

**Appendix Figure A3.**
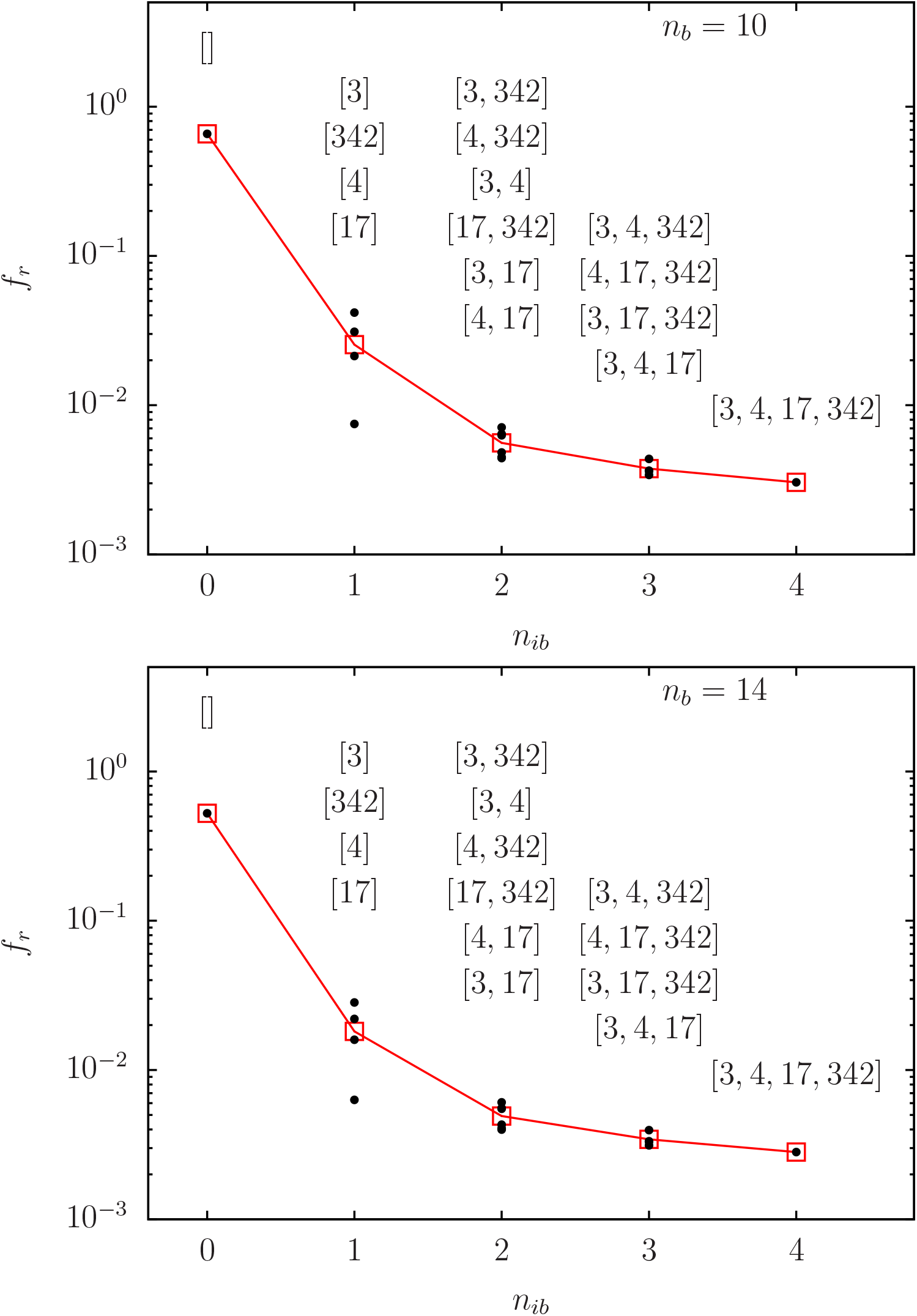
As Figure 6 but for *n*_*b*_ = 10 (top panel) and *n*_*b*_ = 14 (bottom panel).

Appendix Figure A3 shows *f*_*r*_ for optimal configurations at *n*_*b*_ = 10 and *n*_*b*_ = 14 (as Figure 6).

**Appendix Figure A4.**
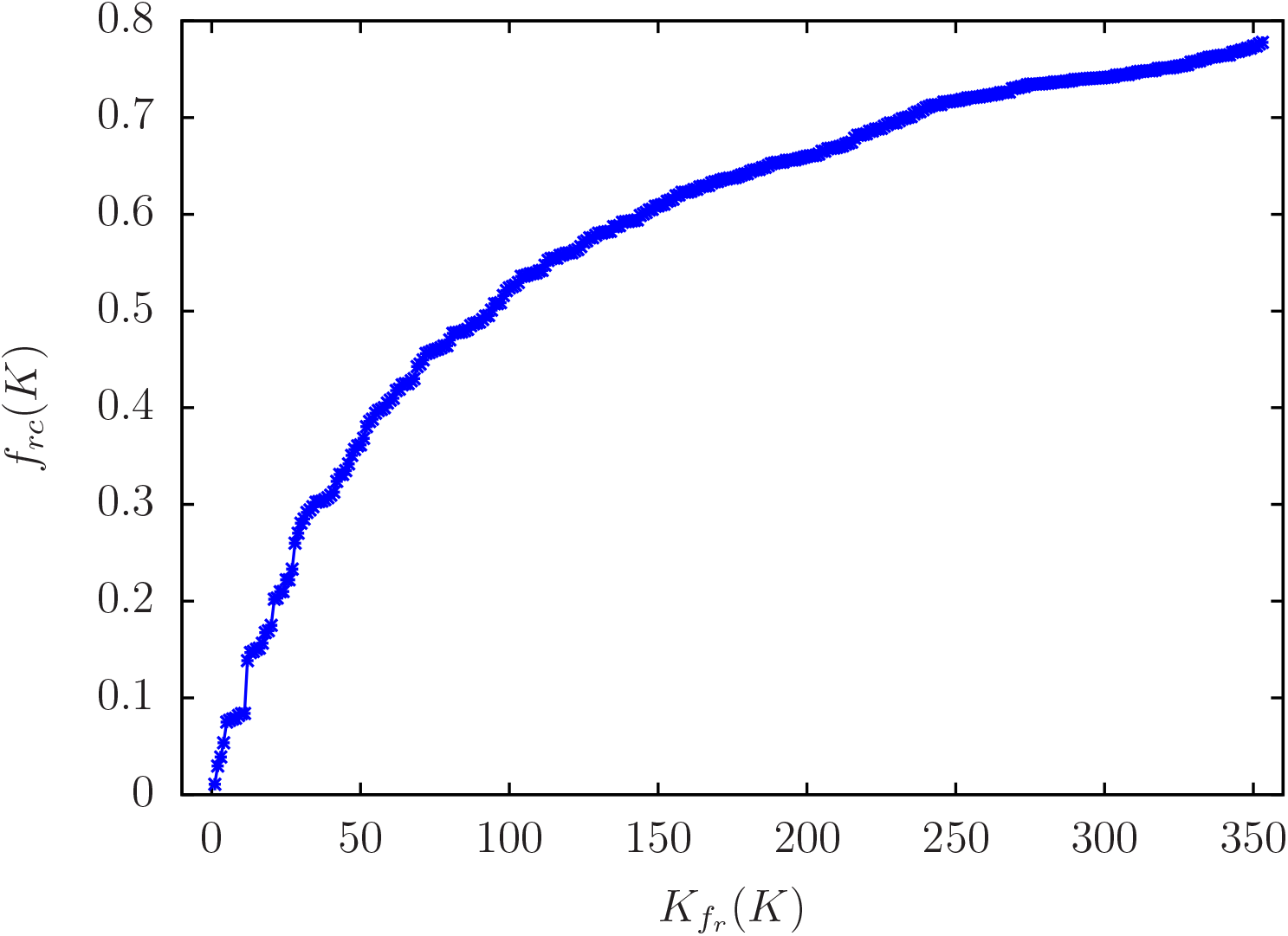
*f*_*rc*_ (*K*) versus 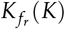 using the data of Figure 8 (i.e. *n*_*b*_ = 6, *f*_*rc*_ (*K*) being computed for *n*_*ib*_ = 1 with one single initial blue node at node *K* belonging to the Erdös group.) The index 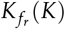 is the index obtained by ordering the *N*_*E*_ = 353 nodes of the Erdös set with increasing values of *f*_*rc*_ (*K*).

Appendix Figure A4 shows the values of *f*_*rc*_(*K*) (for *n*_*b*_ = 6) computed from one initial (variable) blue node *K* belonging to the Erdös group ordered by increasing values of *f*_*rc*_(*K*) inside this group.

**Appendix Figure A5.**
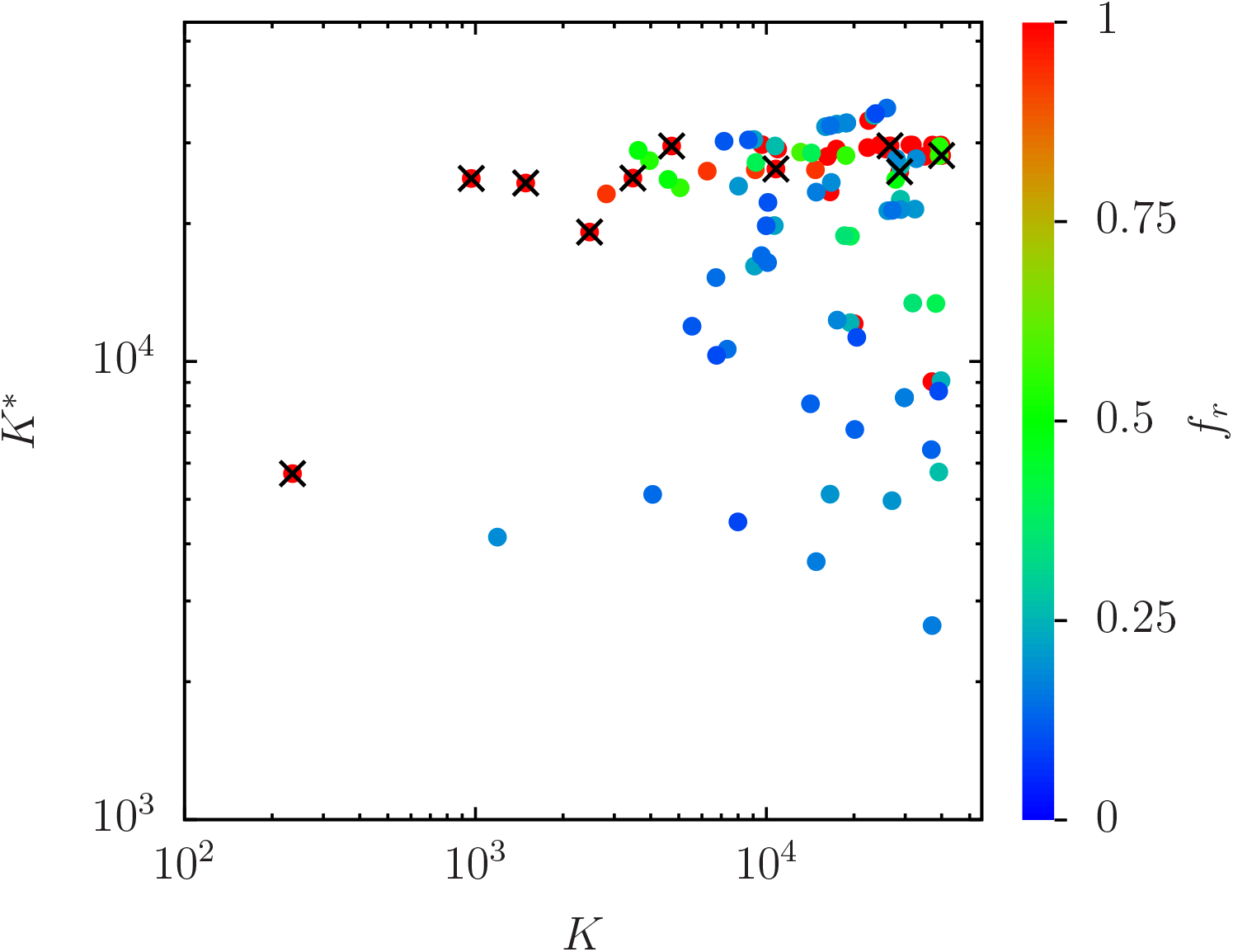
The positions of the 100 nodes with top 100 *f*_*r*_ (*i*) values of the (non network averaged) red outcome for individual nodes *i* in the *K*-*K*^∗^ plane in a zoomed double logarithmic representation (colored full circles) using the data for *n*_*ib*_ = 4 of Figure 6 with the configuration [3, 4, 17, 342] for the initial blue nodes. The color of each data point corresponds to the value of *f*_*r*_ (*i*) according to the color bar. There are 26 nodes *i* with *f*_*r*_ (*i*) = 1 including the 10 permanently fixed red nodes (black cross symbols; *K*_*g*_ = 1, …, 10 in Table 1) and 16 additional nodes and there are 36 (100) nodes *i* with *f*_*r*_ (*i*) *>* 0.5 (*f*_*r*_ (*i*) ≥ 0.08850).

Appendix Figure A5 shows for the data for *n*_*ib*_ = 4 of Figure 6 with the initial blue node configuration [3, 4, 7, 342] the 100 nodes with top individual *f*_*r*_(*i*) values of the red outcome in the *K*-*K*^∗^ plane.

**Appendix Figure A6.**
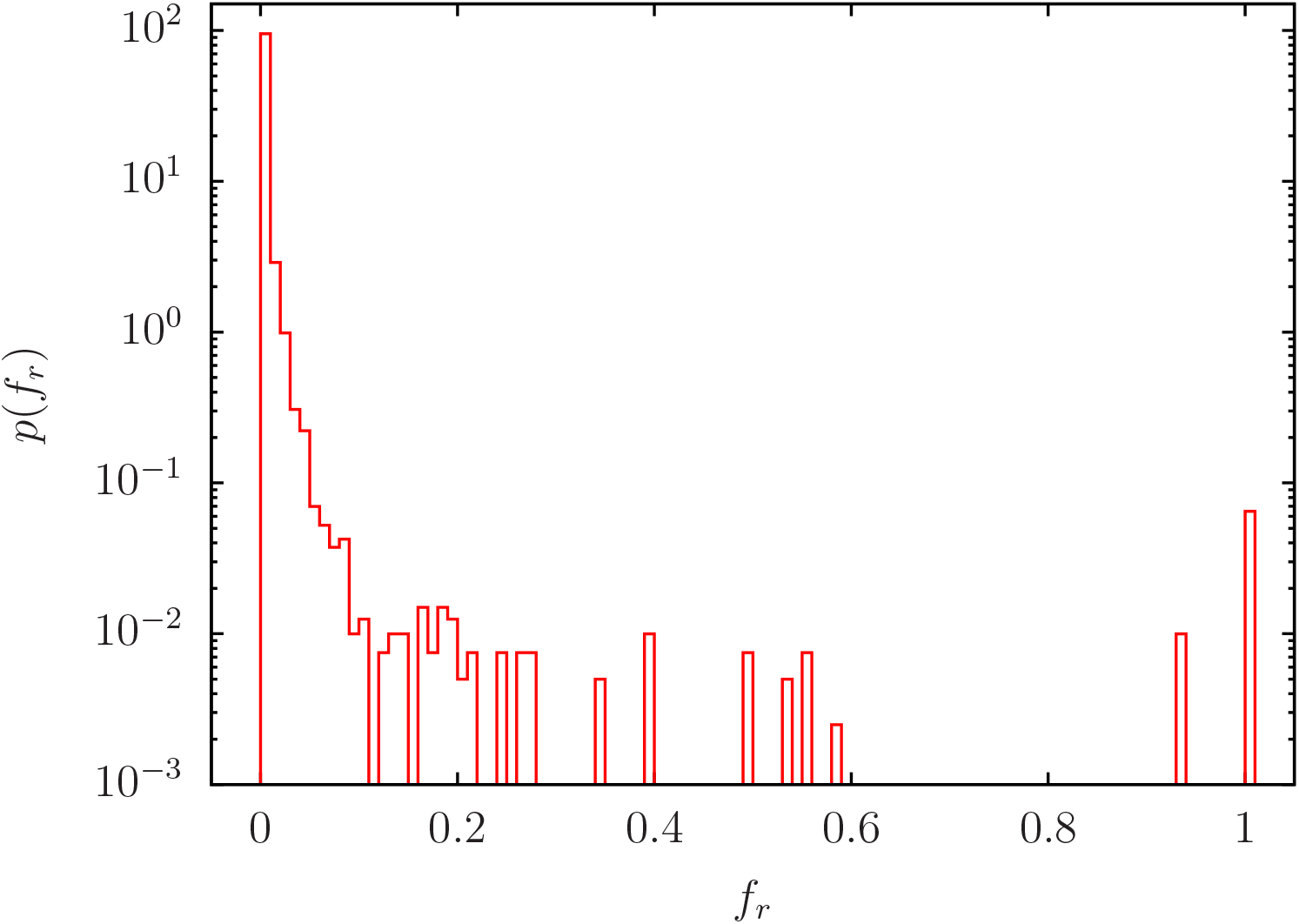
Histogram distribution *p*(*f*_*r*_) for the data of Figure A5 using the values *f*_*r*_ (*i*) for all nodes with bin width Δ *f*_*r*_ = 0.01. The normalization is as in Figure 1 by an integral.

Appendix Figure A6 shows the distribution *p*(*f*_*r*_) for the data of Figure A5. The peak at *f*_*r*_ = 1 corresponds to the 26 nodes *i* with *f*_*r*_(*i*) = 1. The large peak at *f*_*r*_ = 0 corresponds to 95% of probability for 0 ≤ *f*_*r*_ ≤ 0.01.

**Appendix Figure A7.**
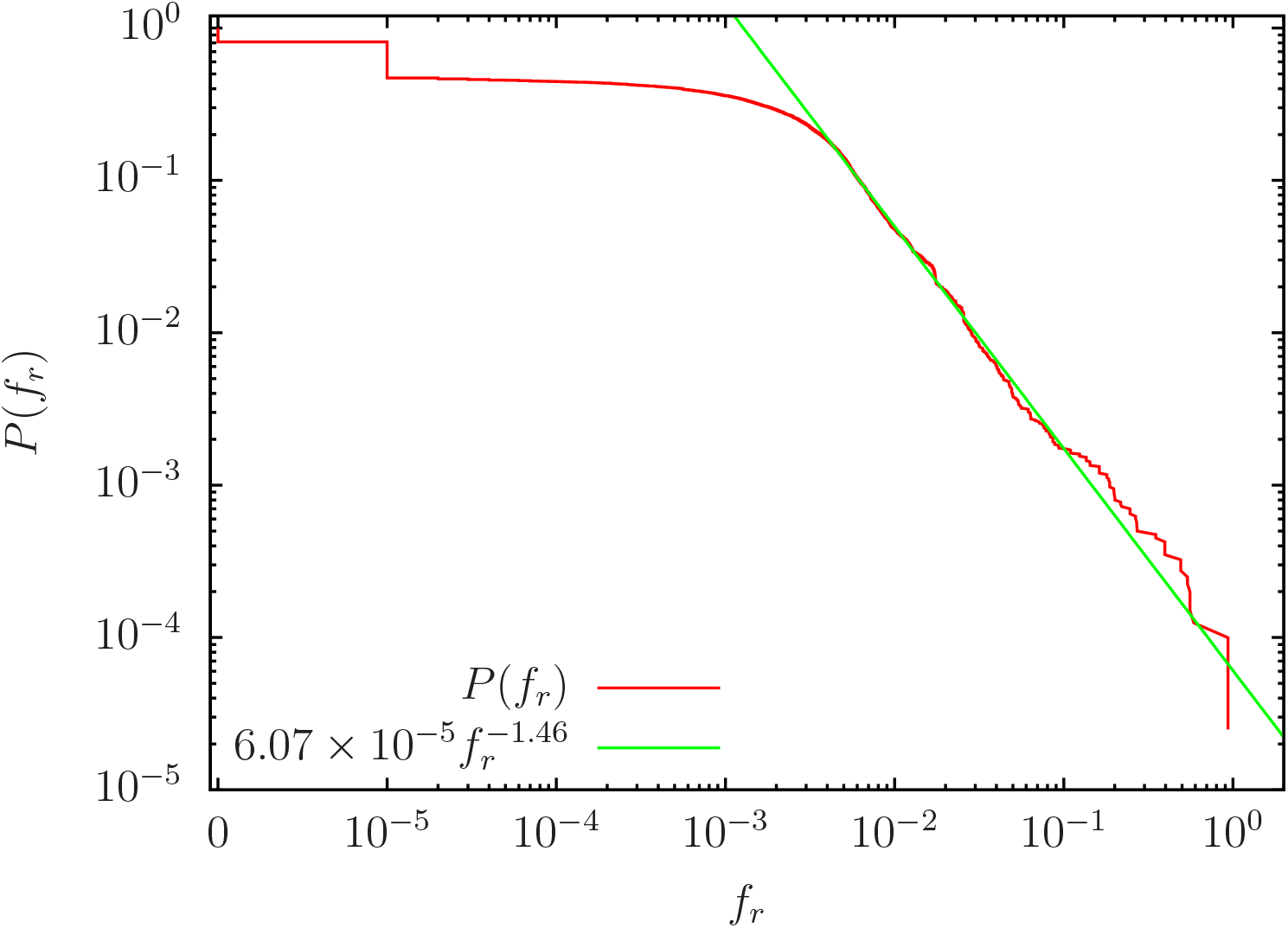
Fraction *P*(*f*_*r*_) of nodes *i* with *f*_*r*_ (*i*) ≥ *f*_*r*_ versus *f*_*r*_ (red curve) for the data of Figs. A5, A6. The presentation is double logarithmic except for the data points at *f*_*r*_ = 0 which have artificially been added at a finite value below 10^−5^ and the 26 nodes *i* with *f*_*r*_ (*i*) = 1 have been excluded in the computation of *P*(*f*_*r*_). The steps/vertical lines at *f*_*i*_ = 0 (*f*_*i*_ = 10^−5^) correspond to 7539 (13696) network nodes *i* having 0 (1) case(s) of red outcome out of the *R* = 100000 random pathway realizations (note that there are 7230 nodes who have always white outcome). The green straight line corresponds to the power law fit 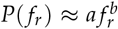 with *a* = (6.07 *±* 0.03) *×* 10^−5^ and *b* = −1.457 *±* 0.001 obtained for the interval *f*^*i*^ ∈ [5 *×* 10^−3^, 4 *×* 10^−2^].

Appendix Figure A7 shows the probability *P*(*f*_*r*_) for a node *i* to have a value *f*_*r*_(*r*) ≥ *f*_*r*_ for the data of Figs. A5, A6. The decay of *P*(*f*_*r*_) is rather well algebraic as 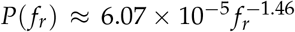 for *f*_*r*_ ≥ 5 *×* 10^−3^ if the 26 nodes *i* with *f*_*r*_(*i*) = 1 are excluded in the computation of *P*(*f*_*r*_). (If these nodes are included the decay is still algebraic in the fit interval *f*_*i*_ ∈ [5 *×* 10^−3^, 4 *×* 10^−2^] but *P*(*f*_*r*_) is significantly above the power law for *f*_*i*_ *>* 4 *×* 10^−2^.) Note that *P*(*f*_*r*_) corresponds to the integrated probability 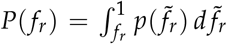 using the probability density *p*(*f*_*r*_) visible in Appendix Figure A6 (apart from the numerical approximation due to the histogram with finite bin width).

## Author Contributions

All authors equally contributed to all stages of this work.

## Acknowledgments

We thank L.Ermann for useful discussions.

